# Structure-guided enhancement of selectivity of chemical probe inhibitors targeting bacterial seryl-tRNA synthetase

**DOI:** 10.1101/586255

**Authors:** Ricky Cain, Ramya Salimraj, Avinash S. Punekar, Dom Bellini, Colin W. G. Fishwick, Lloyd Czaplewski, David J. Scott, Gemma Harris, Christopher G. Dowson, Adrian J. Lloyd, David I. Roper

## Abstract

Aminoacyl-tRNA synthetases are ubiquitous and essential enzymes for protein synthesis and also a variety of other metabolic processes, especially in bacterial species. Bacterial aminoacyl-tRNA synthetases represent attractive and validated targets for antimicrobial drug discovery if issues of prokaryotic versus eukaryotic selectivity and antibiotic resistance generation can be addressed. We have determined high resolution X-ray crystal structures of the *Escherichia coli* and *Staphylococcus aureus* seryl-tRNA synthetases in complex with aminoacyl adenylate analogues and applied a structure-based drug discovery approach to explore and identify a series of small molecule inhibitors that selectively inhibit bacterial seryl-tRNA synthetases with greater than two orders of magnitude compared to their human homologue, demonstrating a route to selective chemical inhibition of these bacterial targets.

## Introduction

The fidelity of protein synthesis is absolutely reliant upon the provision of specific amino acids by aminoacyl-tRNA molecules for use by the ribosome.^1^ Errors in this process cause defects in protein folding and function leading to cell death.^2^ Each of the 20 amino acids has its own aminoacyl-tRNA synthetase (aaRS) which catalyses the attachment of the amino acid to its cognate tRNA. Despite the fact that all aaRSs share the same overall mechanism, it has long been recognised that there is clearly significant diversity between bacterial, mammalian and archaeal enzymes to allow for synthetic and natural product discrimination between pathogen and host enzymes^3-5^. In addition, in some situations, several different amino acids are able to bind to non-cognate aaRSs, requiring an *in vivo* editing function allowing for the possibility of exploiting this feature for future antimicrobial discovery^6^. For example, the amino acid serine is able to bind alanyl-tRNA synthetase (AlaRS) and threonyl-tRNA synthetase (ThrRS) in addition to its cognate seryl-tRNA synthetase (SerRS)^7^. This incorrect binding is rectified in nature by numerous proofreading mechanisms^6, 8^. However, in this context, one of the major challenges presented by aaRS as targets for antimicrobial drug discovery is their ubiquitous presence in organisms and particularly with respect to bacterial infection in human tissues requiring exploration of strategies that allow for bacterial selectivity to prevent issues of specificity and toxicity^9^.

Aminoacyl sulfamoyl adenosines (aaSAs) are non-hydrolysable mimetics of the aminoacyl adenylate intermediate (aaAMP) formed during the aaRS catalytic cycle and are potent inhibitors of these enzymes.^10^ A significant number of natural product inhibitors mimic these reaction intermediates forming tight binding complexes with substantial affinity competing effectively with natural substrates. Of those, mupirocin is the most prominent example that has found clinical utility as a topical treatment for soft tissue infections. Mupirocin targets the IleRS enzyme and utilises a hydrophobic “tail” in addition to an aminoacyl adenylate warhead to bind to its target^11^. By contrast to many antibiotics in clinical use, seryl sulfamoyl adenosine (SerSA, **1**) can bind and inhibit AlaRS and ThrRS in addition to SerRS and hence is a multi-targeting inhibitor^7, 12^. It can be predicted therefore that SerSA would require mutations in several of these enzymes before a resistance phenotype could be conferred.

The protein databank (PDB) contains X-ray crystal structures of SerRS from *Thermus thermophilus*^*13, 14*^, *Methanosarcina barkeri*^*15*^, *Pyrococcus horikoshii*^*16*^, *Candida albicans*^*17*^, *Arabidopsis thaliana*^*18*^, *Methanopyrus kandleri*^*19*^, *Trypanosoma brucei*, human cytoplasmic^20^ and bovine mitochondrial^21^. It is therefore evident that there is a distinct lack of structural data available for clinically relevant bacteria. Although the *Escherichia coli* SerRS^22^ structure was solved in 1990, the coordinates were not deposited in the PDB thereby hampering efforts in antimicrobial structure-based drug discovery (SBDD). Moreover, the X-ray crystal structure of human SerRS in complex with SerSA reveals specific conformational changes upon catalysis necessary for function, which are not found in bacterial homologues providing further perspectives upon differences in structure that may allow prokaryotic from eukaryotic specificity.^20^ In this study we set out to increase the available structural information for human bacterial pathogens and use this to investigate the possibilities for designing bacteria-specific SerRS enzyme inhibitors using a SBDD approach.

## Results

### Crystal structures of SerRS in complex with SerSA inhibitor

The crystal structures of full-length SerRS from *E. coli* (*Ec*SerRS) and *Staphylococcus aureus* (*Sa*SerRS) in complex with the SerSA inhibitor were solved at 1.50 Å and 2.03 Å, respectively (**Fig. 1**, **Supplementary Table 1**). In both structures, the SerSA inhibitor is unambiguously determined by the electron density maps. SerSA is bound deep into a well-conserved SerRS aminoacylation catalytic pocket and stabilized by a network of hydrogen bond interactions from the residues in motif 2, motif 3 and the serine-binding TxE motif (**Fig. 1a**) - a typical binding mode in all class 2 aaRSs. Superimposition of *Ec*SerRS, *Sa*SerRS and human cytoplasmic SerRS (*Hs*SerRS, PDB ID: 4L87^20^) structures show a high degree of similarity as evidenced by the RMSD values (**Supplementary Table 2**). The orientation of the bound SerSA inhibitor is comparable in all three structures. However, the N-terminal tRNA-binding domain (i.e. the two-stranded anti-parallel coiled coil making the long helical arm) protruding away from the active site pockets in the compared structures shows large conformational changes resulting in a high RMSD.^20^ The purine ring of the adenosine in SerSA interacts with a conserved phenylalanine (F287 in *Ec*SerRS, F281 in *Sa*SerRS and F321 in *Hs*SerRS) via a π-π stacking interaction (**Fig. 1b**). The M284 in *Ec*SerRS, L278 in *Sa*SerRS and V318 in *Hs*SerRS are positioned such that they provide main chain hydrogen bond interactions with the ring nitrogens(**Fig. 1c**). The seryl moiety of SerSA extends deep into the pocket to interact with T237, E239, R268, E291 and S391 in *Ec*SerRS and equivalent residues in *Sa*SerRS and *Hs*SerRS. We note the presence of a highly coordinated water molecule 3Å away from the N^3^ of the adenine moiety of the adenylate **(Fig. 1b-c**), a feature that has previously been described in class II synthetase enzymes.^23^ In *Sa*SerRS the octahedral coordination of a magnesium ion (**Fig 1c**) is observed in the active site via Glu349,the SerSA sulfone and water molecules, reminiscent of the magnesium ion observed in both the *Candida albicans* SerRS and *Hs*SerRS.

**Figure 1:**
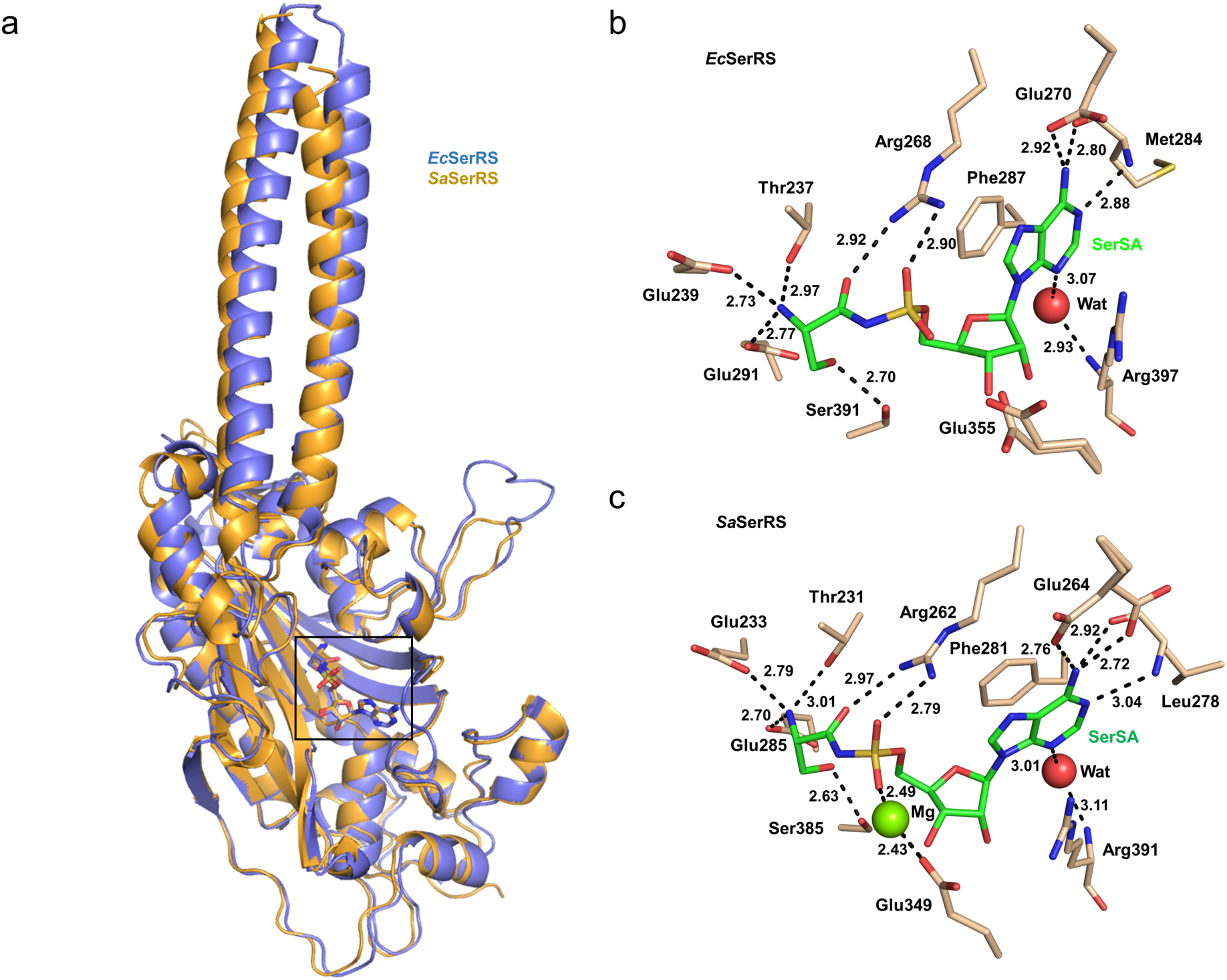
Binding mode of SerSA to *E. coli* and *S. aureus* SerRS. **a**: Superposition of *Ec*SerRS (blue) and *Sa*SerRS (gold) with SerSA bound (boxed). **b**: Interactions of SerSA (green sticks) with *Ec*SerRS chain A. Water represented as a red sphere. Hydrogen bond interactions shown as black dashes. **c**: Interactions of SerSA with *Sa*SerRS.

### Design and synthesis of the selectivity probe

The X-ray crystal structures of *Ec*SerRS, *Sa*SerRS and the *Hs*SerRS (PDB ID: 4L87) were superimposed in Maestro (Schrödinger, LLC).^24^ Interestingly, a thorough analysis of the active site pockets revealed a small extension in the hydrophobic cavity adjacent to the C-2 position of SerSA in the *Ec*SerRS and *Sa*SerRS structures (**1**). This pocket is centred around a glycine at positions 396 and 390 in *Ec*SerRS, *Sa*SerRS respectively. This hydrophobic cavity extension is absent in the *Hs*SerRS (**Supplementary Fig. 1b**) as it is filled by the bulkier side-chain of threonine at position 434. The conserved nature of this structural difference is reflected in amino acid sequence alignments of selected Gram-positive and Gram-negative bacterial pathogens when compared to cytoplasmic and mitochondrial variants of the human, bovine and mouse SerRS enzymes (**Supplementary Fig. 2a**). Moreover, inspection of an alignment of the Gram-positive and Gram-negative bacterial pathogens shows this glycine to be part of a 12 amino acid region of conservation ending in an arginine (397 in *Ec*SerRS, 391 in *Sa*SerRS) suggestive of an invariant bacterial structural feature absent in eukaryotic homologues. (**Supplementary Fig. 2b**). Exploiting such a conserved feature for antimicrobial drug discovery extends the range of bacteria that can potentially be targeted whilst also reducing the chances of mutation-induced drug resistance.

A focused structure-activity relationship (SAR) series with variants at the C-2 position of SerSA adenosine was designed to investigate the steric tolerance of the hydrophobic cavity and to establish the degree of selectivity for the bacterial SerRS over the *Hs*SerRS (**Fig. 2a**). *In silico* molecular docking of the designed selectivity probes into the active site pockets of the *Ec*SerRS, *Sa*SerRS and *Hs*SerRS crystal structures (**Supplementary Methods**) and visual analysis of the predicted docking poses (**Supplementary Fig. 1c-d**) suggested that chloro-and iodo-seryl sulfamoyl adenylate derivatives **2** and **3** respectively would not achieve selectivity since **2** and **3** were predicted to interact equally as well with both the bacterial and *Hs*SerRS. Compounds **4**-**8** were however predicted to exhibit selectivity for the bacterial SerRS over the *Hs*SerRS.

**Figure 2:**
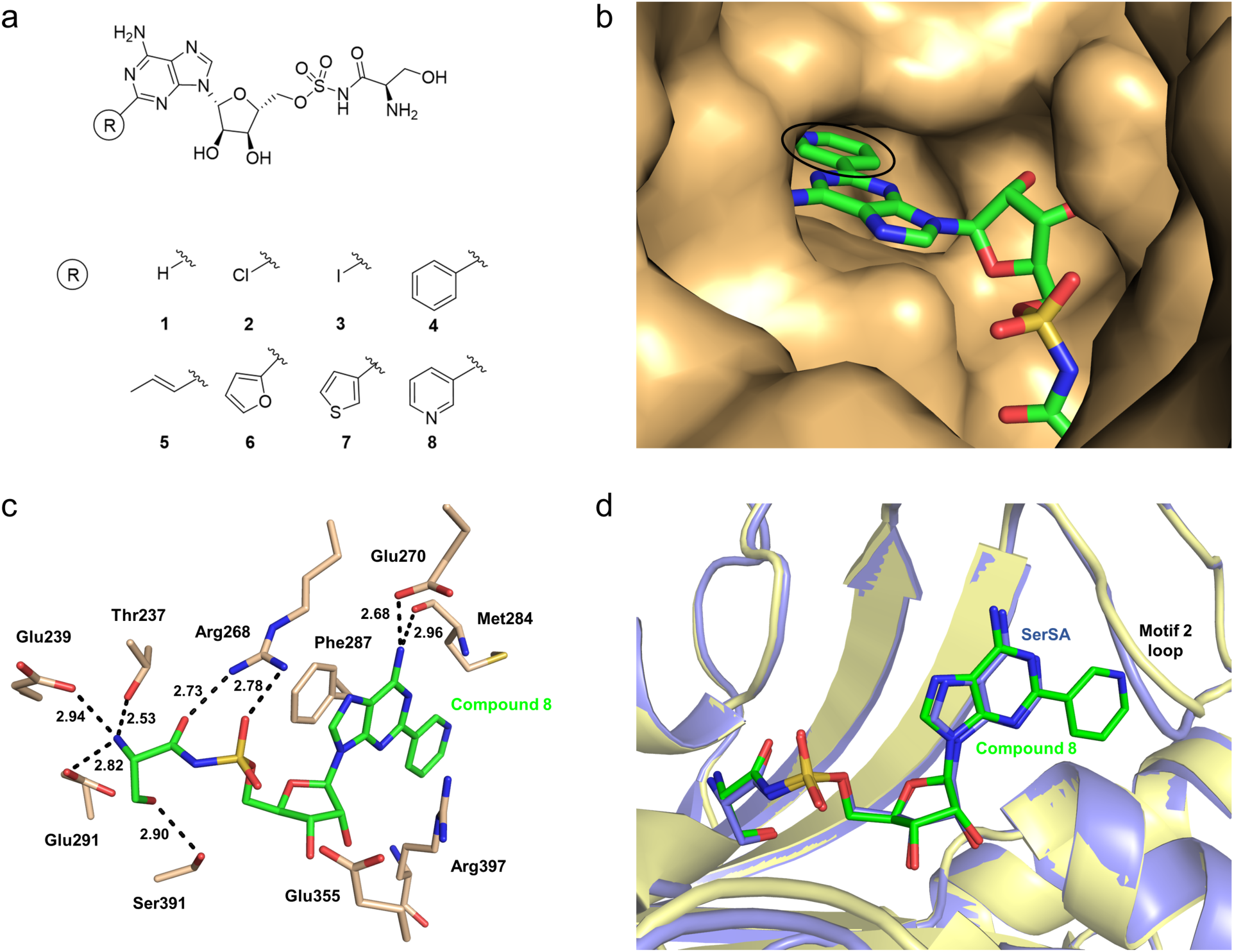
Comparison of binding of SerSA and compound 8 to *Ec*SerRS active site. a: The chemical structures of the compounds used in this study. **b**: Pyridyl group of compound 8 (circled) positioned in the active site. **c**: Interactions of compound 8 (green sticks) with *Ec*SerRS. Hydrogen bond interactions are shown as black dashes. d: Superposition of *Ec*SerRS:SerSA (blue) with *Ec*SerRS:compound **8**.

The bulkier groups located at the 2 position of compounds **4**-**8** were predicted to be accommodated in the pocket of the bacterial enzymes. However, due to the steric hindrance from the T434 residue in the *Hs*SerRS, compounds **4-8** were predicted to change the torsional angle between the adenine and ribose sugar upon binding to the *Hs*SerRS. As a result of the torsional change the π-π stacking interactions with F287 and the backbone interaction to V318 are lost leading to a weaker predicted binding affinity and therefore increased selectivity for the bacterial SerRS (**Supplementary Fig. 1d**).

Preparation of SerSA selectivity probes was initiated by the acid-catalysed protection of the commercially available 2-chloroadenosine or 2-iodoadenosine (Fluorochem, UK) to provide the acetyl-protected adenosines (95-97%) (**Supplementary Synthesis**). A Suzuki coupling reaction between the protected adenosine and desired boronic acid species (20-70%) was conducted,^25^ before sulfonation using sulfonyl chloride to afford the sulfonamide (90-95%). The sulfonamide was then coupled to the succinimide activated protected serine **(S1, Supplementary Information)** to yield the protected product (40-50%). Removal of the benzyl group was accomplished by treatment with a solution of boron trichloride dimethyl sulfide complex (2M in DCM),^26^ and the resulting alcohol was treated with trifluoroacetic acid and water to yield compounds **2-8** (**2**-**8**, see **Experimental Section** and **Supplementary Information** for details).

### Bacterial SerRS inhibition by selectivity probe

Using a continuous spectrophotometric assay that specifically measures the adenylate formation reaction^7^, compounds **2-8** were evaluated for the inhibition of ATP-dependent aminoacyl adenylate formation by *Ec*SerRS and *Sa*SerRS enzymes and compared their half maximal inhibitory concentration (IC_50_) values with the parent SerSA, compound **1** (**Supplementary Figure 3 and Supplementary Table 4**). Compounds **2-8** were active against *Ec*SerRS and *Sa*SerRS with IC_50_ values ranging from 378 nM to 52.7 µM (**Table 1**). Compound **2** exhibited sub-micromolar inhibition of the *Sa*SerRS and *Ec*SerRS with IC_50_s of 262 nM and 445 nM respectively. Compound **3** also exhibited sub-micromolar inhibition of *Sa*SerRS with an IC_50_ of 378 mM but weaker inhibition of *Ec*SerRS with an IC_50_ of 1.36 µM. Compounds **4**-**8** all manifested low micromolar inhibition against both bacterial SerRS (**Table 1**). A general trend is observed where increasing the size of the group at the 2 position of the adenylate decreases the binding affinity to the bacterial synthetase. Alanyl sulfamoyl adenosine (AlaSA, **9**) and threonyl sulfamoyl adenosine (ThrSA **10**) were also evaluated for inhibition against *Ec*SerRS and *Sa*SerRS (**Supplementary Table 5**). AlaSA **9** showed no inhibitory activity against either enzyme at 1 mM while ThrSA **10** manifested IC_50_s of 285 µM and 231 µM against *Ec*SerRS and *Sa*SerRS respectively, thus exhibiting much weaker binding than the designed selectivity probes. These results highlight the key role of the beta-hydroxyl of the serine to the overall binding of the compound within the adenylate formation site in these enzymes and the overall inhibitory properties of seryl adenylate inhibitors modified around the C-2 position of the SerSA adenosine.

**Table 1:** IC_50_ values of designed chemical probes against seryl-tRNA synthetases. Assays were conducted as reported.^7^.

| No. | X | IC<sub>50</sub> *EcSerRS* (μM) | IC<sub>50</sub> *SaSerRS* (μM) | IC<sub>50</sub> *HsSerRS* (μM) |
| --- | --- | --- | --- | --- |
| **1 (SerSA)** | H | 0.21 ± 0.03 | 0.23 ± 0.49 | 2.17 ± 0.21 |
| **2** | Cl | 0.45 ± 0.05 | 0.26 ± 0.03 | 67.3 ± 4.67 |
| **3** | I | 1.36 ± 0.12 | 0.38 ± 0.04 | 24.0 ± 2.26 |
| **4** | C<sub>6</sub>H<sub>5</sub> | 17.7 ± 1.42 | 52.7 ± 4.81 | >1000 ± >100 |
| **5** | *trans*-Propenyl | 9.38 ± 0.70 | 3.46 ± 0.47 | >1000 ± >100 |
| **6** | 2-Furyl | 36.2 ± 2.41 | 32.4 ± 3.56 | >1000 ± >100 |
| **7** | 3-Thienyl | 1.44 ± 0.09 | 1.24 ± 0.12 | >1000 ± >100 (ppt) |
| **8** | 3-Pyridyl | 6.65 ± 0.64 | 6.34 ± 0.71 | >1000 ± >100 (ppt) |

SerRS, Seryl t-RNA synthetase. (ppt) precipitation observed at 1000 μM. Errors were calculated as s.d. of at least three independent measurements.

### *Hs*SerRS inhibition by selectivity probes

Measurement of the IC_50_ inhibition kinetics of the original SerSA, compound **1** against the bacterial and human SerRS enzymes, reveals a 10-fold difference, in favour of greater specificity of the inhibitor for the bacterial enzymes. Compounds **2-8** were subsequently screened for inhibition of the *Hs*SerRS (**Table 1**) using the same assay system. Assay measurements of compounds **2** and **3**, revealed a 31-fold and 11-fold increase in IC_50_ against the bacterial SerRS and *Hs*SerRS, indicating that compounds **2** and **3** did not exhibit selectivity overall and had lower affinity than SerSA **1**. Overall the observed IC_50_ of compounds **2-8** increased with respect to the parental adenylate **1** but remarkably, inhibition of the *Hs*SerRS was effectively abolished in compounds **4**-**8** with IC_50_ values greater than 1 mM, revealing significant selectivity of these compounds towards the tested bacterial SerRS. The best of these compounds (**7**), with a 3-Thienyl at the C-2 position of the SerSA adenosine had an increase in IC_50_ over SerSA **1** of 6.8 and 8.4 fold for *Ec*SerRS and *Sa*SerRS respectively, with effectively negligible binding to the *Hs*SerRS. The observed selectivity overall was attributed to the increased size of **4-8** making them unable to fit into the hydrophobic pocket located in the human cytoplasmic SerRS active site due to the presence of T434 as previously hypothesised.

### Binding studies of SerSA and compound 8 to *Ec*SerRS

To independently measure the binding characteristics of the original adenylate SerSA and the derivatives synthesised in this study, we measured binding affinity using iso isothermal titration calorimetry (ITC). The binding stoichiometry and affinity of SerSA **1** and compound **8** to *Ec*SerRS was determined using ITC because compound **8** had the best solubility of the synthesised compounds. Titration of SerSA to *Ec*SerRS resulted in a steep slope in the binding isotherm suggesting a very tight binding of the inhibitor to the enzyme. Interestingly, fitting of this binding isotherm using a single site model showed a 2:1 SerSA:SerRS stoichiometry with an overall dissociation constant K_d_ = 1.27 nM (**Supplementary Fig. 5a**). The combination of very high affinity and low enthalpy unfortunately prevented an accurate measurement of K_d_ for SerSA at the individual binding sites.

By contrast, titration of compound **8** to *Ec*SerRS resulted in a binding isotherm (2:1 compound **8**:SerRS stoichiometry) that after fitting using a two independent sites model clearly showed two distinct binding sites with dissociation constants K_d1_ = 0.29 µM and K_d2_ = 1.92 µM (**Supplementary Fig. 5b**). As K_d2_ > 4 K_d1_, there is apparent mild negative cooperativity within the system. In both experiments, a negative enthalpy value detected for such a tight interaction indicates the role of hydrogen bond and electrostatic interactions in the stabilisation of the enzyme–inhibitor complex. The observation of two binding sites for SerSA and compound 8 prompted us to investigate the oligomeric state of the *Ec*SerRS in solution, which are typically dimers in solution.^27^ Analytical ultracentrifugation (AUC) experiments were carried out with *Ec*SerRS to confirm the oligomeric state of the protein in the presence and absence of SerSA and compound **8 (Supplementary Table 6)**. The results confirmed that *Ec*SerRS, both with and without inhibitors, appeared with a molecular weight that is consistent with a dimer in solution (**Supplementary Fig. 4**). The observed SerSA and compound **8** binding stoichiometry is consistent with the previous structural findings showing two SerSA molecules bound to two distinct sites in the X-ray crystal structure of *Candida albicans* SerRS (PDB ID: 3QO8)^28^. In this structure the second SerSA binding site is located 26 Å distant from the active site and appears to play no role in the enzyme function or protein-protein interaction as described by the authors ^28^.

### Structural basis of selectivity probe binding to *Ec*SerRS

To understand the molecular basis of the selectivity probe towards bacterial SerRS, we attempted a series of co-crystallisation screening of *Ec*SerRS and *Sa*SerRS. Despite extensive screening, we were unable to find hit conditions to co-crystallise *Sa*SerRS in the presence of compound **7** or **8**. However, we were successful in obtaining crystals of *Ec*SerRS amenable to soaking with compound **8**, which yielded a 2.6 Å resolution structure (**Fig. 2a-d**). The *Ec*SerRS-SerSA complex structure was solved in the space group P1 containing two monomers that associate tightly to form a dimer. In contrast, the *Ec*SerRS-compound **8** complex structure was solved in space group P6_1_22 with 1 molecule in the asymmetric unit. Compound **8** binds in a similar fashion to SerSA in *Ec*SerRS making key interactions with the residues in motif 2, motif 3 and the serine-binding TxE motif as described above (**Fig. 2c**). The 3-pyridyl group of compound **8** snuggly fits into the hydrophobic cavity without any other obvious interactions (**Fig. 2b**) with movement of the motif 2 loop observed to accommodate the pyridyl group (**Fig. 2d**).

We analysed both structures for presence of a second adenylate-binding site as found in the *Candida albicans* SerRS-SerSA structure (PDB ID: 3QO8)^28^. No density for the second adenylate was found in the *Ec*SerRS-SerSA structure, but density for two additional ligand molecules were observed in the *Ec*SerRS-compound **8** structure (**Fig. 3a**). These ligands were found to bind away from the active site in positions distinct to that observed in the *Candida albicans* SerSA (**Fig. 3b**). Electron density for the complete compound was observed for the ligand in the active site (**Fig. 3c**) and a second ligand which π-π -stacks with a third ligand from a symmetry-related molecule for which electron density is only observed for its purine and pyridyl rings (**Fig. 3d**). The seryl moiety of this third ligand is likely to not make any interactions with the protein and be flexible due to the absence of electron density for this region of the compound. As such the presence of this third ligand molecule is likely to be a crystallographic artefact of the high concentration of compound used for soaking and this structure provides evidence for a potential second binding site that is supported by the ITC data.

**Figure 3:**
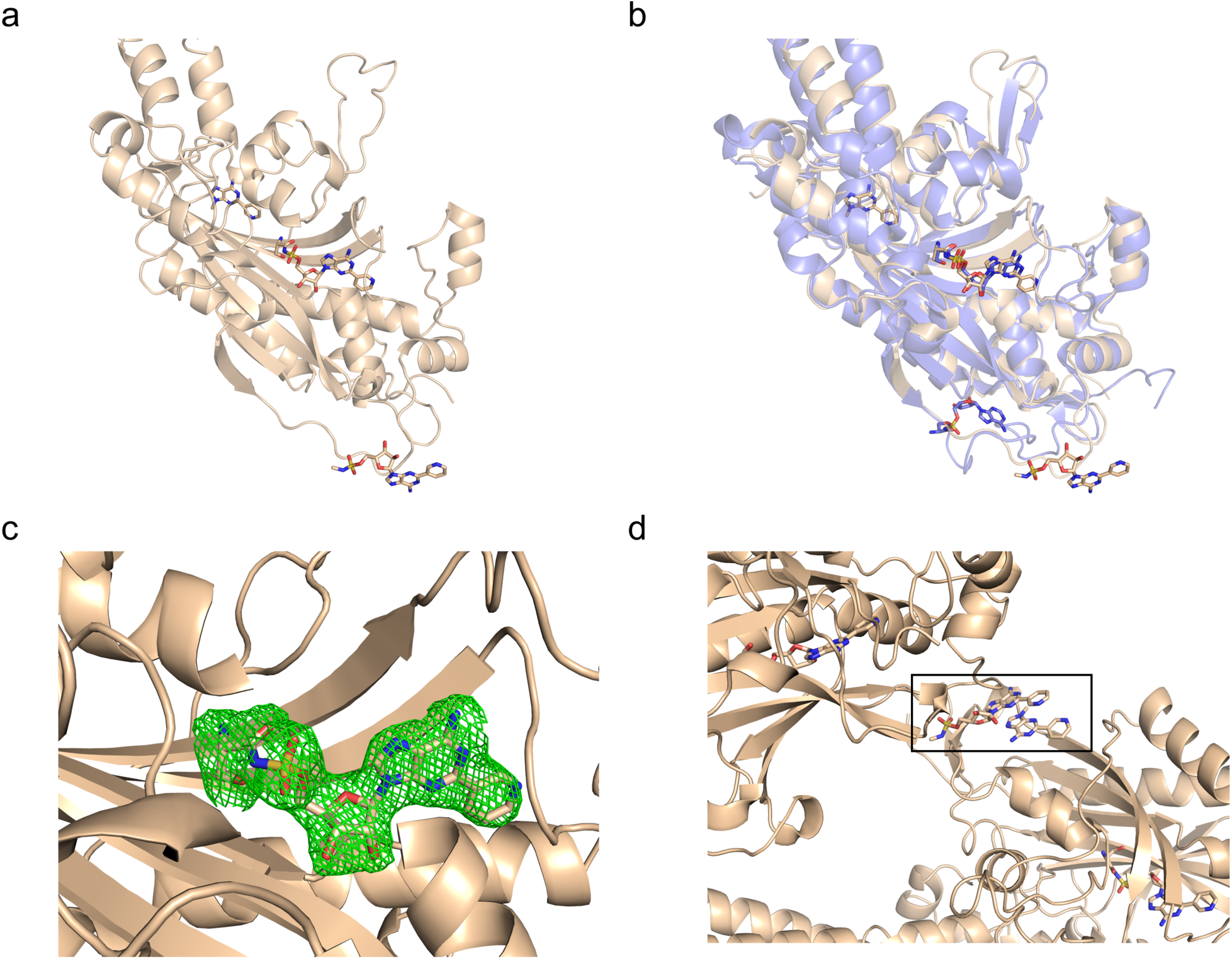
Second binding site of compound 8 to *Ec*SerRS. **a**: Binding positions of compound **8** to *Ec*SerRS. **b**: Overlay of *Ec*SerRS:compound **8** (wheat) with *Candida albicans* SerRS (blue) **c**: Overlay of F_o_-F_c_ omit map of compound **8** in *Ec*SerRS active site contoured at σ 3. **d**: Interaction of two molecules of compound **8** (boxed) from *Ec*SerRS symmetry-related molecules.

## Discussion

In summary, we demonstrate the use of structure-based drug design to identify selective inhibitors of exemplar SerRS enzymes from Gram-positive and Gram-negative pathogens on the WHO list of bacteria for which new antibiotics are urgently needed. Previous studies have investigated inhibiting protein synthesis via inhibition of specific aaRS activities leading to the identification of a number of potent antibiotics which have progressed through into clinical studies^29-32^. Rapid development of resistance to these synthetase inhibitors has halted their clinical evaluation^33^. The reported alternative approach herein has been a proof of principle example of the capability of structure-based drug design in modifying a multi-targeting aaRS inhibitor to achieve selectivity.

We have analysed the compounds produced in this study using the bioinformatics tool Entryway (www.entry-way.org), which classifies molecules that are likely to be capable of accumulating in Gram-negative bacteria. Whilst the compounds fulfil the requirements for globularity and contain the required primary amine the number of rotatable bonds exceeds the limits normally founds in antimicrobials.^34^ Further work is required to achieve clinically viable compounds that can permeate the cell membrane but the crystal structures here, nonetheless, provide a foundation for structure-based drug design of novel selective inhibitors which multi-target the bacterial aminoacyl-tRNA synthetases.

## Methods

### Synthesis

Full experimental details and characterisation of the compounds are given in Supplementary information.

### Protein expression and purification

*Ec*SerRS (from *E. coli* strain B ER2560) and *Sa*SerRS (from *S. aureus seg*50 (1150)) were cloned into pET52b(+) vector (Merck Millipore, Germany) using the NcoI and SacI restriction sites allowing for the production of protein with a thrombin cleavable C-terminal His_10_-tag. *Ec*SerRS and *Sa*SerRS were overexpressed in Lemo21(DE3) cells grown in Auto Induction Media – Terrific Broth (Formedium) supplemented with 100 µg/ml ampicillin at 37°C for 8 hours followed by overnight growth at 25°C. Cells were harvested by centrifugation at 5000 rpm in a JLA 8.1000 rotor (Beckman Coulter) for 15 min, and the pellet was re-suspended in buffer A (50 mM Tris-HCl pH 7.5, 500 mM NaCl, 30 mM Imidazole). The cells were disrupted by sonication at 70 % amplitude for 30 sec on ice and 8 pulses. The lysate was centrifuged at 18,000 rpm in a JA 25.50 rotor (Beckman Coulter) for 30 mins. The supernatant was decanted, passed through a 0.2 µm filter and applied to a 5 ml HisTrap column (GE healthcare, USA). The bound protein was eluted with a gradient of buffer B (50 mM Tris-HCl pH 7.5, 500 mM NaCl, 500 mM Imidazole) (0-100% over 50 ml) on an ÄKTA Pure (GE healthcare, USA) at 2 ml/min. The protein was dialyzed into 2 L of buffer A with thrombin cleavage (1 unit/µg). The protein was passed through the 5 ml HisTrap column to remove the cleaved His-tag and other contaminants. The proteins typically present over 95 % purity at this stage as judged via SDS-PAGE gel and were taken for crystallization trials after dialysis into 20 mM Tris-HCl pH 7.5, 200 mM NaCl and 1 mM MgCl_2_. Further purification was used for protein used for kinetic and binding studies to ensure complete removal of thrombin using a HiLoad 16/600 Superdex 200 pg column (GE Healthcare, USA) in 20 mM Tris-HCl pH 7.5, 200 mM NaCl and 1 mM MgCl_2_. The purified protein was subsequently stored in 50 % glycerol at -80°C. *HsSerRS* was expressed and purified as previously described.^35^

### Crystallisation and structure solution

Co-crystals of *Ec*SerRS in the presence of SerSA were obtained from a drop set up in 96-well sitting drop format with 20 mg ml ^−1^ protein and ten-fold molar excess of SerSA. Drops consisted of 100 nl protein preincubated with SerSA and 100 nl reservoir solution with a reservoir volume of 95 µl. Crystals were obtained from a drop containing 0.2 M sodium phosphate monobasic monohydrate, pH 4.7 and 20 % (w/v) PEG 3350 following incubation at 4°C and cryoprotected in reservoir solution containing 25 % (v/v) ethylene glycol.

Co-crystals of His-tagged *Sa*SerRS were obtained from a drop set up with 20 mg ml^−1^ protein in the presence of ten-fold molar excess of SerSA in 24-well hanging drop format. Drops consisted of 1 µl protein preincubated with SerSA and 1 µl reservoir solution with a reservoir volume of 500 µl. Plates were incubated at 4°C and crystals obtained in 0.2 M sodium malonate pH 5.0 and 13 % (w/v) PEG 3350. Crystals were cryoprotected for 10 s in reservoir solution containing 20 % (v/v) ethylene glycol and ten-fold molar excess of SerSA.

Crystals of apo-*Ec*SerRS were obtained at 21°C from a 24-well hanging drop format as described above with 30 mg ml^−1^ protein in a crystallisation condition consisting of 0.1 M sodium citrate pH 5.5, 0.8 M lithium sulfate and 0.05 M ammonium sulfate. A single crystal was soaked for 30 mins in 0.1 M sodium citrate pH 5.5, 0.75 M lithium sulfate, 0.05 M ammonium sulfate, 20 % (v/v) ethylene glycol and 100 mM compound **8** (10 % (v/v) DMSO in final solution).

All crystals were flash frozen in liquid nitrogen and diffraction data collected at 100 K at beamlines I03 and I04 (Diamond Light Source, United Kingdom). Data was indexed and integrated using iMosflm^36^ and scaled using Aimless in CCP4^37^ or autoPROC^38^ was used in the DLS auto-processing pipeline. The crystal structure of aq_298 (PDB 2DQ3, unpublished) was used as a search model in Phaser MR^39^ to solve the structures of *Ec*SerRS and *Sa*SerRS by molecular replacement. Phenix^40^ and Buster^41^ were used for iterative rounds of refinement with model building carried out in COOT.^42^ Figures were made using PyMOL (Schrödinger, LLC).

### Kinetic analyses

SerRS assays were performed at 37°C in a Cary 100 UV/Vis double beam spectrophotometer with a thermostatted 6X6 cell changer. The final assay volume was 0.2 ml, containing 50 mM HEPES adjusted to pH 7.6, 10 mM MgCl_2_, 50 mM KCl, 1 mM dithiothreitol, 10% (v/v) dimethylsulphoxide, 10 mM D-glucose, 0.5 mM NADP+, 1.7 mM.min yeast hexokinase and 0.85 mM.min *L. mesenteroides* glucose 6-phosphate dehydrogenase. Concentrations of SerRS, amino acid, adenylate (Ap4A) and pyrophosphate were as stated in the text. Unless otherwise stated, background rates were acquired in the absence of amino acid, which was then added to initiate the full reaction. Assays were continuously monitored at 340 nm, to detect reduction of NADP+ to NADPH, where ΔNADPH; 340nm = 6220 M^−1^ cm^−1^ (**Supplementary Table 3**). Kinetic constants relating to substrate dependencies and IC_50_ values for inhibitors were extracted by non-linear regression using GraphPad Prizm 7.00.

### Isothermal titration calorimetry

Calorimetric titrations of *Ec*SerRS with SerSA and/or compound **8** were performed on a VP-ITC microcalorimeter (MicroCal) at 25°C and measured in triplicates. The gel-filtration purified *Ec*SerRS was concentrated and dialysed overnight against the ITC buffer (20 mM Tris-HCl, pH 7.5 and 200 mM NaCl) at 4°C. All the solutions were degassed by sonication. The overnight dialysis ITC buffer was used to prepare SerSA and compound **8** solutions. The *Ec*SerRS (3 µM for SerSA and 7 µM for compound **8**) in the sample cell (1.445 ml) was titrated with ligand solution (70 µM of SerSA and 140 µM of compound **8**) in the syringe (280 ul). The *Ec*SerRS -SerSA ITC experiments consisted of a preliminary 2 µl injection followed by 52 successive 5 µl injections. The *Ec*SerRS -compound **8** ITC experiments consisted of a preliminary 2 µl injection followed by 26 successive 10 µl injections. Each injection lasted 20 s with an interval of 120 s between consecutive injections. The solution in the reaction cell was stirred at 307 rpm throughout the experiments. The heat response data for the preliminary injection was discarded and the rest of the data was used to generate binding isotherm. The data were fit using either the one binding site model or the two independent binding sites model included in the Origin 7.0 (MicroCal). Thermodynamic parameters, including association constant (K_a_), enthalpy (ΔH), entropy (ΔS) and binding stoichiometry (N) were calculated by iterative curve fitting of the binding isotherms. The Gibbs free energy was calculated using ΔG = ΔH -TΔS.

### Analytical ultracentrifugation

All experiments were performed at 50000 rpm, using a Beckman Optima analytical ultracentrifuge with an An-50Ti rotor. Data were recorded using the absorbance (at 280 nm with 10 µm resolution and recording scans every 20 seconds) and interference (recording scans every 60 seconds) optical detection systems. The density and viscosity of the buffer was measured experimentally using a DMA 5000M densitometer equipped with a Lovis 200ME viscometer module. The partial specific volume for the protein constructs were calculated using Sednterp from the amino acid sequences. For characterisation of the protein samples, SV scans were recorded for a dilution series, starting from 0.8 mg/mL. Where a ligand was included, this was present at 400 uM (a 20-fold excess over the highest concentration protein sample). Data were processed using SEDFIT, fitting to the c(s) model.^43^ Figures were made using GUSSI.^44^.

### Data availability

The crystallographic data that support the findings of this study are available from the Protein Data Bank (http://www.rcsb.org). *Ec*SerRS:SerSA, 6R1M; *Sa*SerRS:SerSA, 6R1N; *Ec*SerRS:compound **8**, 6R1O. Additional data that support the findings of this study are available from the corresponding author upon reasonable request.

## Acknowledgements

We acknowledge funding from Medical Research Council (MRC) for Innovation Grant MR/M017893/1 and BBSRC Flexible Talent Mobility Account award BB/R506588/1 that supported this work directly, BBSRC BB/R013411/1for AUC supporting infrastructure at the Harwell Campus and EPSRC AMR bridging the gaps grant EP/M027503/1 that provided initial pump priming activity. We thank Diamond Light Source for beam time (proposal MX14692) and staff at beamlines I03 and I04 for their assistance. We would like to acknowledge Dr Michael Haertlein of the Institut Laue Langevin, Grenoble, France for supply of the *HsSerRS* expression construct used in this study and Dr Yann-Vaï Le Bihan of the Institute of Cancer Research, London, United Kingdom for advice and support during the structure refinement process. We acknowledge undergraduate students Archna Shah and Sarah Brocklesby for initial cloning of the bacterial synthetase genes.

## Author contributions

A.J.L, C.G.D, and D.I.R conceived the research. R.C with the help of C.W.G.F and L.C designed, synthesized and characterized compounds used for the study. R.S produced and purified the enzymes used in biochemical studies. R.S and D.B crystallised the seryl adenylate inhibitors with *Ec*SerRS and *Sa*SerRS and collected X-ray data. R.S and A.S.P solved and refined the final crystal structures as presented. R.C with the help of A.J.L carried out the kinetic studies. R.C, R.S, A.S.P with the help of D.J.S conducted and analysed the results of ITC binding assays. D.J.S and G.M carried out AUC analysis. R.C, R.S and D.I.R wrote the manuscript. All authors discussed the results and contributed to the final manuscript.

## Supplementary Information

Any supplementary information, chemical compound information and source data are available in the online version of the paper.

