## Supplementary Fig for "Structure-guided enhancement of selectivity of chemical probe inhibitors targeting bacterial seryl-tRNA synthetase"

### A) List of Figures and Schemes

***Supplementary Figure 1***: Binding modes of designed seryl sulfamoyl adenylate selectivity probes. (a) 3D spatial representation of the seryl sulfamoyl adenylate derivatives. (b) Structural overlay of *S. aureus* SerRS (Yellow, PDB ID 6R1N) and human cytoplasmic SerRS (Grey, PDB ID: 4L87) active site showing the key residue change near the 2 position of the sulfamoyl adenylate inhibitor from Gly390 in the bacterial form to Thr434 in the human form. (c) Predicted binding modes of seryl sulfamoyl adenylates to *E. coli* SerRS (PDB ID 6R1M) using AutoDock 4.2. (d) Predicted binding modes of seryl sulfamoyl adenylates to human cytoplasmic SerRS using AutoDock 4.2.

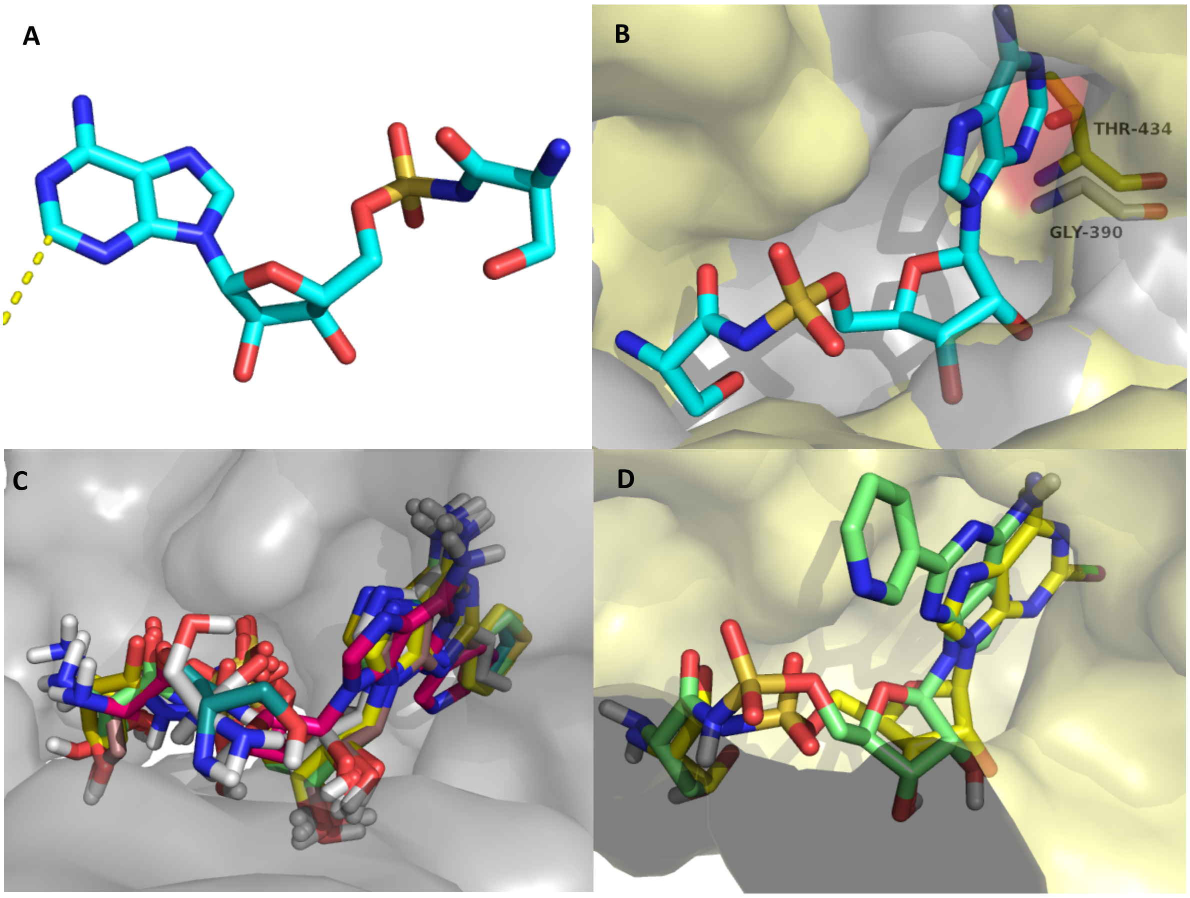

***Supplementary Figure 2a:*** Sequence alignment prepared by EMBL-EBI MUSCLE[1] of representative Gram-positive and Gram-negative bacterial pathogens against cytoplasmic and mitochondrial variants of the human, bovine and mouse SerRS enzymes. The blue boxed feature indicates the conserved glycine (396 in *Ec*SerRS) found in the bacterial enzymes which is replaced by threonine (434 in the cytoplasmic *Hs*SerRS) and proline (458 in the mitochondrial *Hs*SerRS) in the cytoplasmic and mitochondrial eukaryotic enzymes respectively. A consensus sequence across all SerRS enzymes in this region is indicated.

```
Homo sapiens(cyto)          KVEFVHMLNATMCATTRTICAILENYQTEKG-ITV
Homo sapiens(mito)          ELQFAHTVNATACAVPRLLIALLESNQQKDGSVLV
Bos taurus(cyto)            KVEFVHMLNATMCATTRTICAILENYQTEKG-ILV
Bos taurus(mito)            ELQFAHTVNATGCAVPRLLIALLESYQQKDGSVLV
Mus musculus(cyto)          KVEFVHMLNATMCATTRTICAILENYQAEKG-IAV
Mus musculus(mito)          ELQFAHTVNATACAVPRVLIALLESNQQKDGSVLV

Escherichia coli            KTRLVHTLNGSGLAVGRTLVAVMENYQQADGRIEV
Acinetobacter baumannii     KTELVHTLNGSGLAVGRTLLAVMENYQREDGSIEI
Pseudomonas aeruginosa      KPELVHTLNGSGLAVGRTLVAVLENYQQADGSIRV
Haemophilus influenzae      KTRLVHTLNGSGLAVGRTLVAVLENYQNADGSITV
Neisseria gonorrhoeae       KNRLVHTLNGSGLAVGRTLVAVLENHQNADGSINI

Staphylococcus aureus       KPELAHTLNGSGLAVGRTFAAIVENYQNEDGTVTI
Streptococcus pneumoniae    KVKLLHTLNGSGLAVGRTVAAILENYQNEDGSVTI
Enterococcus faecium        KVQYTHTLNGSGLAVGRTVTAILENYQNEDGSVTI
Clostridium difficile       KAEYVHTLNGSGLAIGRCLAAILENYQQADGSVVV
Listeria monocytogenes      KPEYVHTLNGSGLALGRTVAAILENYQDADGSVRI
                            :      :*.:  *   * . *::*. *  .* : :
                                    NxxxxAxxR
```

***Supplementary Figure 2b:*** Sequence alignment prepared by EMBL-EBI MUSCLE[1] of representative Gram-positive and Gram-negative bacterial pathogens. The blue box indicate a 12 amino acid sequence within the adenylate binding pocket. The glycine indicated by the arrow at the 11th position in this sequence is replaced by threonine and proline in the cytoplasmic and mitochondrial eukaryotic enzymes as described in Supplementary Figure 2a. As described in the main text the glycine is at position 396 in *Ec*SerRS.

```
Escherichia coli            SSCSNVWDFQARRMQARCRSKSDKKTRLVHTLNGSGLAVGRTLVAVMENYQQADGRIEVP
Acinetobacter baumannii     SSCSNMGDFQARRMKARFRMD-QKKTELVHTLNGSGLAVGRTLLAVMENYQREDGSIEIP
Pseudomonas aeruginosa      SSCSNCGDFQARRMQARYRNPETGKPELVHTLNGSGLAVGRTLVAVLENYQQADGSIRVP
Haemophilus influenzae      SSCSNMWDFQARRMQARCKAKGDKKTRLVHTLNGSGLAVGRTLVAVLENYQNADGSITVP
Neisseria gonorrhoeae       SSCSNCEDFQARRMKARFKDE-NGKNRLVHTLNGSGLAVGRTLVAVLENHQNADGSINIP

Staphylococcus aureus       SSCSNCTDFQARRANIRFKRDKAAKPELAHTLNGSGLAVGRTFAAIVENYQNEDGTVTIP
Streptococcus pneumoniae    SSCSNTEDFQARRAQIRYRDEADGKVKLLHTLNGSGLAVGRTVAAILENYQNEDGSVTIP
Enterococcus faecium        SSCSNCEDFQARRAMIRYRNE-EGKVQYTHTLNGSGLAVGRTVTAILENYQNEDGSVTIP
Clostridium difficile       SSCSNFEDFQARRAGIRFKRDKKSKAEYVHTLNGSGLAIGRCLAAILENYQQADGSVVVP
Listeria monocytogenes      SSCSNFESFQARRANIRFRREPGSKPEYVHTLNGSGLALGRTVAAILENYQDADGSVRIP
                            *****   *****   * .     *    ************ . *::****. ***: :*
```

***Supplementary Figure 3:*** Dependence of *E. coli* SerRS, *S. aureus* SerRS and human cytoplasmic SerRS on a) [Pyrophosphate], b) [Serine], and c) [$Ap_4A$].

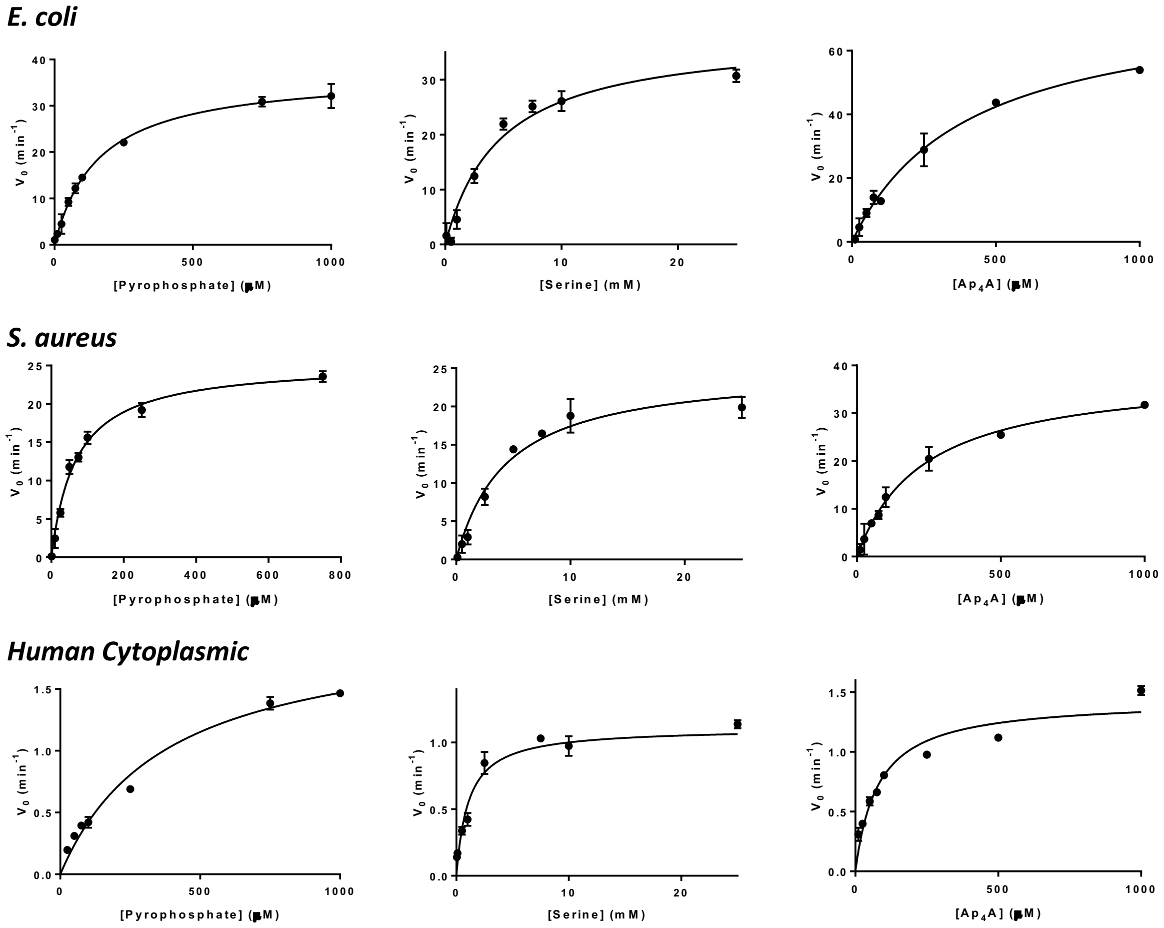

***Supplementary Figure 4:*** **Sedimentation coefficient distributions of EC SerRS. A:** Absorbance data **B:** Interference data **C:** Absorbance data **D:** Interference data **E:** Interference data.

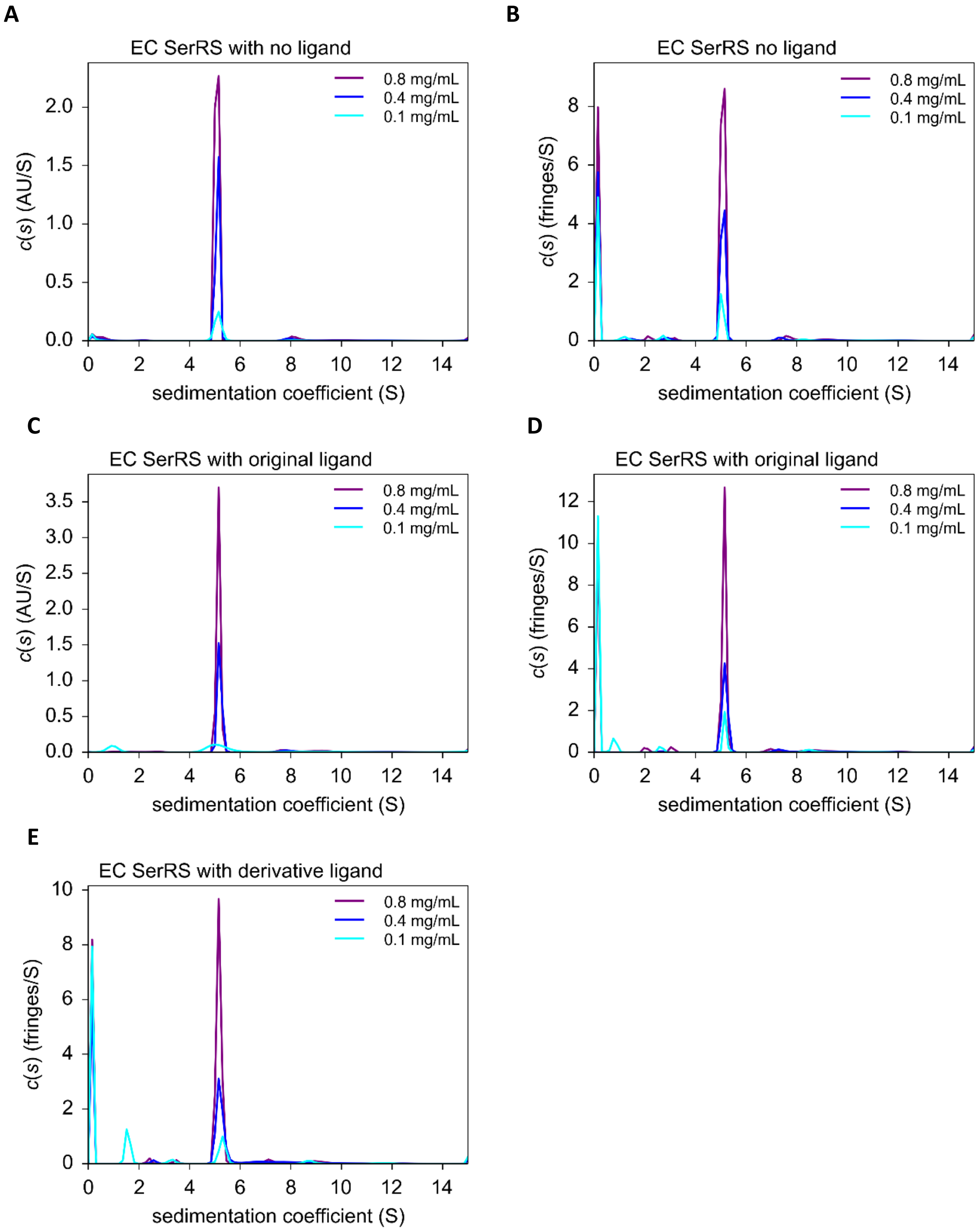

***Supplementary Figure 5:*** **Isothermal Titration Calorimetry (ITC) curves. A:** Titration of 70 µM SerSA **1** into 3 µM EC SerRS **B:** Titration of 140 µM compound **8** into 7 µM EC SerRS.

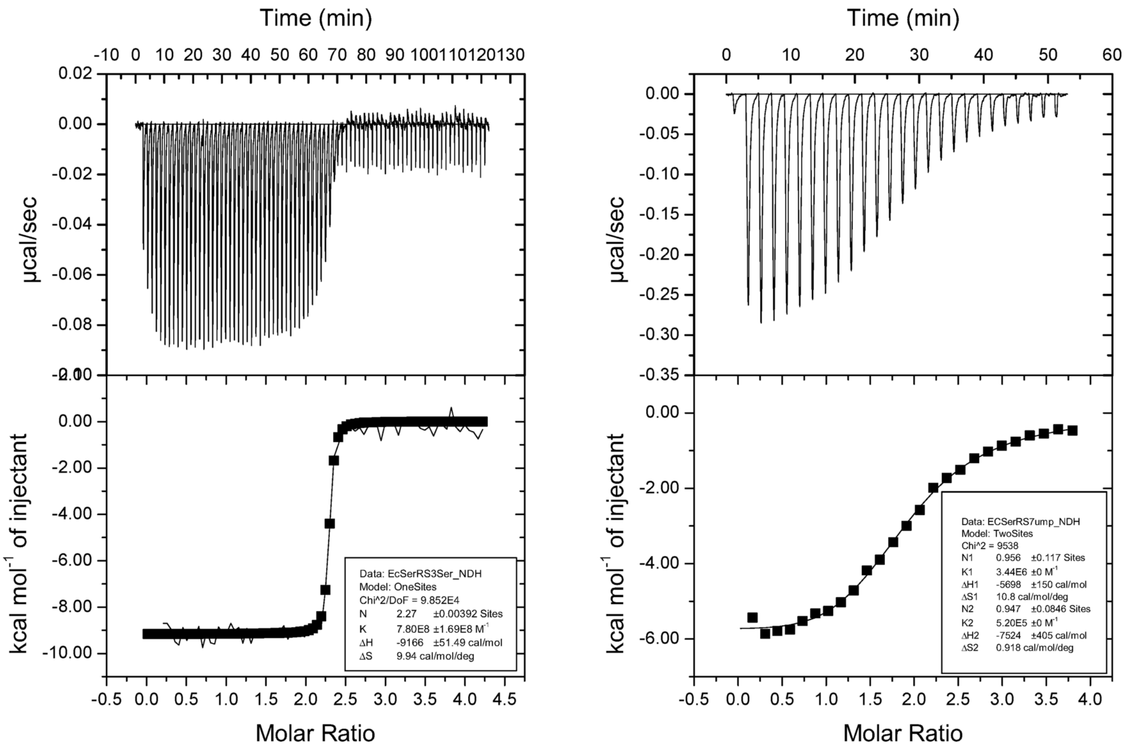

#### Supplementary Tables

***Supplementary Table 1*:** Data collection and refinement statistics of SerSA **1** bound bacterial SerRS structures and **8** bound to *E. coli* SerRS

| Data Collection | *E. coli* SerRS: SerSA | *S. aureus* SerRS: SerSA | *E. coli* SerRS: compound 8 |
|---|---|---|---|
| X-ray source | DLS (I03) | DLS (I04) | DLS (I04) |
| Wavelength | 0.9762 | 0.9795 | 0.9795 |
| Space Group | *P*1 | $C222_1$ | $P6_122$ |
| Cell Dimensions | | | |
| *a*, *b*, *c* (Å) | 63.07, 63.52, 75.89 | 94.44 116.41 91.61 | 85.41, 85.41, 317.72 |
| α, β, γ | 75.44, 70.25, 89.20 | 90, 90, 90 | 90, 90, 120 |
| Molecules/ asymmetric unit | 2 | 1 | 1 |
| Resolution (Å) | 61.29 - 1.50 (1.53 - 1.5) | 91.61 - 2.03 (2.06 - 2.03) | 158.86 - 2.60 (2.64 - 2.60) |
| Redundancy | 2.4 (2.5) | 11.8 (7.7) | 28.5 (30.3) |
| $R_{merge}$ | 0.048 (0.269) | 0.106 (0.807) | 0.179 (1.876) |
| $CC_{1/2}$ | 0.997 (0.829) | 0.999 (0.856) | 0.999 (0.949) |
| $I/\sigma$ | 9.3 (2.5) | 16.1 (2.3) | 17.4 (2.0) |
| Completeness (%) | 95.5 (93.8) | 99.8 (97.7) | 100 (100) |
| **Refinement** | | | |
| Resolution (Å) | 59.18 - 1.50 | 57.25 - 2.03 | 27.96 - 2.60 |
| No. of reflections | 163577 | 33037 | 22215 |
| $R_{work}/R_{free}$ | 0.162/0.180 | 0.173/0.221 | 0.182/0.215 |
| Atoms | | | |
| Protein | 6831 | 3332 | 3310 |
| Ligand | 58 | 29 | 82 |
| Solvent | 1000 | 416 | 192 |
| B factor (Å$^2$) | | | |
| Protein | 24.18 | 35.51 | 63.01 |
| Ligand | 15.04 | 23.12 | 62.37 |
| Solvent | 38.63 | 45.79 | 70.74 |
| RMSD | | | |
| Bonds (Å) | 0.010 | 0.010 | 0.010 |
| Angles (°) | 1.03 | 1.05 | 1.09 |
| Ramachandran (%) | | | |
| Favoured | 99 | 99 | 97 |
| Allowed | 1 | 1 | 3 |
| Outlier | 0 | 0 | 0 |

***Supplementary Table 2*****:** RMSD scores generated using PDBeFOLD[2] when comparing the full-length SerRS structures from *E. coli* (PDB ID: 6R1M), *S. aureus* (PDB ID: 6R1N), and human (PDB ID: 4L87). *E. coli* SerRS (chain A) was used as a reference for superposing other structures. Number of matched residues shown in parentheses.

| | ***Ec*****SerRS (chain A)** | ***Ec*****SerRS (chain B)** | ***Sa*****SerRS** | ***Hs*****SerRS** |
|---|---|---|---|---|
| ***Ec*****SerRS (chain A)** | - | 1.18 (426) | 1.37 (410) | 1.66 (407) |

a. Lower score is better

***Supplementary Table 3*****:** Components of the phosphate exchange assay

| **Reagent** | ***Sa*****SerRS Final concentrations** | ***Ec*****SerRS Final Concentrations** | ***Hs*****SerRS Final Concentrations** |
|---|---|---|---|
| 0.25 M HEPES, 50 mM $MgCl_2$, adjusted to pH 7.6 | 50 mM HEPES, 10 mM $MgCl_2$ | 50 mM HEPES, 10 mM $MgCl_2$ | 50 mM HEPES, 10 mM $MgCl_2$ |
| 0.1 M DTT | 1 mM | 1 mM | 1 mM |
| 2 M KCl | 50 mM | 50 mM | 50 mM |
| 1 M D-Glucose | 10 mM | 10 mM | 10 mM |
| 10 mM $NADP^+$ | 0.5 mM | 0.5 mM | 0.5 mM |
| Coupling Enzymes (Leuconostoc mesenteroides hexokinase & glucose 6-Phosphate dehydrogenase) | 1.7 and 0.85 µM/min/mL respectively | 1.7 and 0.85 µM/min/mL respectively | 1.7 and 0.85 µM/min/mL respectively |
| DMSO (+inhibitor) | 10 % (v/v) | 10% (v/v) | 10% (v/v) |
| Enzyme (aaRS) | 0.343 µM | 0.284 µM | 7.62 µM |
| 10 mM PPi | 1mM | 1 mM | 1 mM |
| 1M Serine | 25 mM | 25 mM | 25 mM |
| AP4A | 250 µM | 250 µM | 250 µM |
| Triton X-100 (10% v/v) | 2 µl | 2 µl | 2 µl |
| Water | 85 µl | 85 µl | 40 µl |

***Supplementary Table 4*****:** Km and Kcat Values

| **AaRS** | **[AaRS]$_{assay}$ (µM)** | **nonvaried substrate** | **[nonvaried substrate] (mM)** | **varied substrate** | **$K_m^{app}$ (µM)** | **$K_{cat}^{app}$ ($min^{-1}$)** | **$K_{cat}^{app}/K_m^{app}$ ($min^{-1}$ $µM^{-1}$)** | **goodness of fit ($r^2$)** |
|---|---|---|---|---|---|---|---|---|
| *Ec*SerRS | 0.284 | L-serine | 25.0 | L-serine | 4706 ± 706 | 38.2 ± 2.10 | 0.00812 | 0.964 |
| | 0.284 | $Ap_4A$ | 0.250 | Ap4A | 402 ± 43.1 | 76.5 ± 3.71 | 0.190 | 0.984 |
| | 0.284 | PPi | 1.00 | PPi | 159 ± 10.2 | 37.2 ± 0.810 | 0.234 | 0.990 |
| *Sa*SerRS | 0.343 | L-serine | 25.0 | L-serine | 4390 ± 654 | 25.1 ± 1.33 | 0.00572 | 0.963 |
| | 0.343 | $Ap_4A$ | 0.250 | Ap4A | 231 ± 23.5 | 38.6 ± 1.53 | 0.167 | 0.979 |
| | 0.343 | PPi | 1.00 | PPi | 68.7 ± 5.30 | 25.4 ± 0.648 | 0.370 | 0.986 |
| *Hs*SerRS | 7.62 | L-serine | 25.0 | L-serine | 1083 ± 156 | 1.11 ± 0.0334 | 0.00102 | 0.947 |
| | 7.62 | $Ap_4A$ | 0.250 | Ap4A | 79.5 ± 11.2 | 1.49 ± 0.0620 | 0.0187 | 0.916 |
| | 7.62 | PPi | 1.00 | PPi | 379 ± 43.6 | 2.03 ± 0.0963 | 0.00535 | 0.980 |

***Supplementary Table 5*: Enzymatic screening results for AlaSA 9 and ThrSA 10.** Assays were conducted as reported.[3]

| No. | Structure | $IC_{50}$ *Ec*SerRS (μM) | $IC_{50}$ *Sa*SerRS (μM) | $IC_{50}$ *Hs*SerRS (μM) |
|---|---|---|---|---|
| **9 (Ala)** | 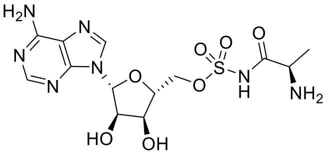 | >1000 ± >100 | >1000 ± >100 | >1000 ± >100 |
| **10 (Thr)** | 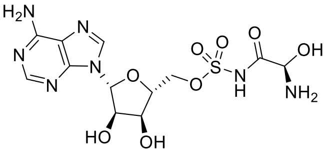 | 285 ± 29.3 | 231 ± 19.0 | >1000 ± >100 |

***Supplementary Table 6*: Species estimated molecular weights from the c(s) analysis.** For each sample concentration the signal-weighted sedimentation co-efficient and the estimated molecular weight of each species is shown, together with the best-fit frictional ratio for the distribution.

| Monomer MW (kDa) | Detection method | Concentration (mg/mL) | Major species | | $f/f_0$ |
|---|---|---|---|---|---|
| | | | MW (kDa) | Sed. Co (S) | |
| **EC SerRS with no ligand** | | | | | |
| 49.2 | Absorbance | 0.8 | 97.5 | 5.08 | 1.40 |
| | | 0.4 | 96.9 | 5.11 | 1.39 |
| | | 0.1 | 93.8 | 5.13 | 1.36 |
| | Interference | 0.8 | 102 | 5.08 | 1.45 |
| | | 0.4 | 104 | 5.06 | 1.46 |
| | | 0.1 | 76.4 | 5.05 | 1.20 |
| **EC SerRS with original ligand** | | | | | |
| 49.2 | Absorbance | 0.8 | 100 | 5.15 | 1.41 |
| | | 0.4 | 92.6 | 5.19 | 1.33 |
| | | 0.1 | 67.8 | 5.16 | 1.08# |
| | Interference | 0.8 | 105 | 5.14 | 1.46 |
| | | 0.4 | 99.9 | 5.16 | 1.41 |
| | | 0.1 | 70.5 | 5.16 | 1.11 |
| **EC SerRS with 8** | | | | | |
| 49.2 | Absorbance | 0.8 | Absorbance too high | | - |
| | | 0.4 | | | - |
| | | 0.1 | | | - |
| | Interference | 0.8 | 103 | 5.16 | 1.43 |
| | | 0.4 | 96.7 | 5.22 | 1.36 |
| | | 0.1 | 41.9 | 5.30 | 0.77# |

### Very poor data fit

#### Supplementary Methods

Chemicals were from commonly used suppliers and used without further purification. Solvents (including dry solvents) for chemical transformations, work-up and chromatography were from Sigma-Aldrich (Dorset, UK) at HPLC grade, and used without further distillation. Silica gel 60 F254 analytical thin layer chromatography (TLC) plates were from Merck (Darmstadt, Germany) and visualized under UV light and/or with potassium permanganate stain. Chromatographic purifications were performed using Merck Geduran 60 silica (40-63 μm) or prepacked SNAP columns on a Biotage Isolera Purification system (Uppsala, Sweden). Deuterated solvents were from Sigma-Aldrich, Chambridge Isotopes and Apollo Scientific Ltd. All $^{1}H$ and $^{13}C$ NMR spectra were recorded using a Bruker Avance 500 MHz spectrometer. All chemical shifts are in ppm relative to the solvent peak, and coupling constants (J) are reported in Hz to the nearest 0.5. High Resolution (HR) mass spectrometry data (m/z) were obtained using a Bruker MaXis Impact instrument with an ESI source and Time of Flight (TOF) analyzer. Fourier transform Infrared (FT-IR) spectra were recorded on a Bruker Alpha Platinum instrument. Melting points were obtained from a Reichert Hot Stage melting point apparatus. HPLC analysis was run on an Agilent 1290 Infinity system equipped with a Supelco Ascentis Express 2.7 μM C18 column (50 x 2.1 mm) using a gradient of 95% solvent A → 95% solvent B (solvent A: $H_2O$ containing 0.1% formic acid; solvent B: 100% MeCN containing 0.1% formic acid), flow rate = 0.5 mL/min and UV detection at 254 nm. Specific rotation measurements were recorded using a Schmidt and Haensch Polartronic H532 polarimeter, using a 100 mm cell and the Sodium D line (589 nm). $[\alpha]_D$ are reported in units of $10^{-1}$ deg $dm^2g^{-1}$.

#### AutoDock Docking protocol

The "*in silico*" docking studies were conducted using AutoDock 4.2[4] as described in the steps below:

1. The crystal structure of the target protein *Ec*SerRS was derived from the study as described in the main text (PDB ID: 6R1M)
2. The active site was defined using maestro.[5] A 20 Å spherical 'cut' of the protein surrounding the active site was designated as the protein receptor.
3. Each ligand was constructed in maestro.[5] The resulting structures were then fully energy minimised using the multiple minimisation tool (MM).
4. Each compound was then docked into the active site of the target synthetase using AutoDock 4.2 which utilises a Lamarckian genetic algorithm. Twenty dockings were conducted for each compound.
5. The resulting docking 'poses' were scored using the AutoDock scoring function.
6. The process was repeated for each synthetase (*Sa*SerRS (PDB ID: 6R1N) and Human cytoplasmic SerRS (4L87))

### Supplementary synthesis

#### Synthetic routes

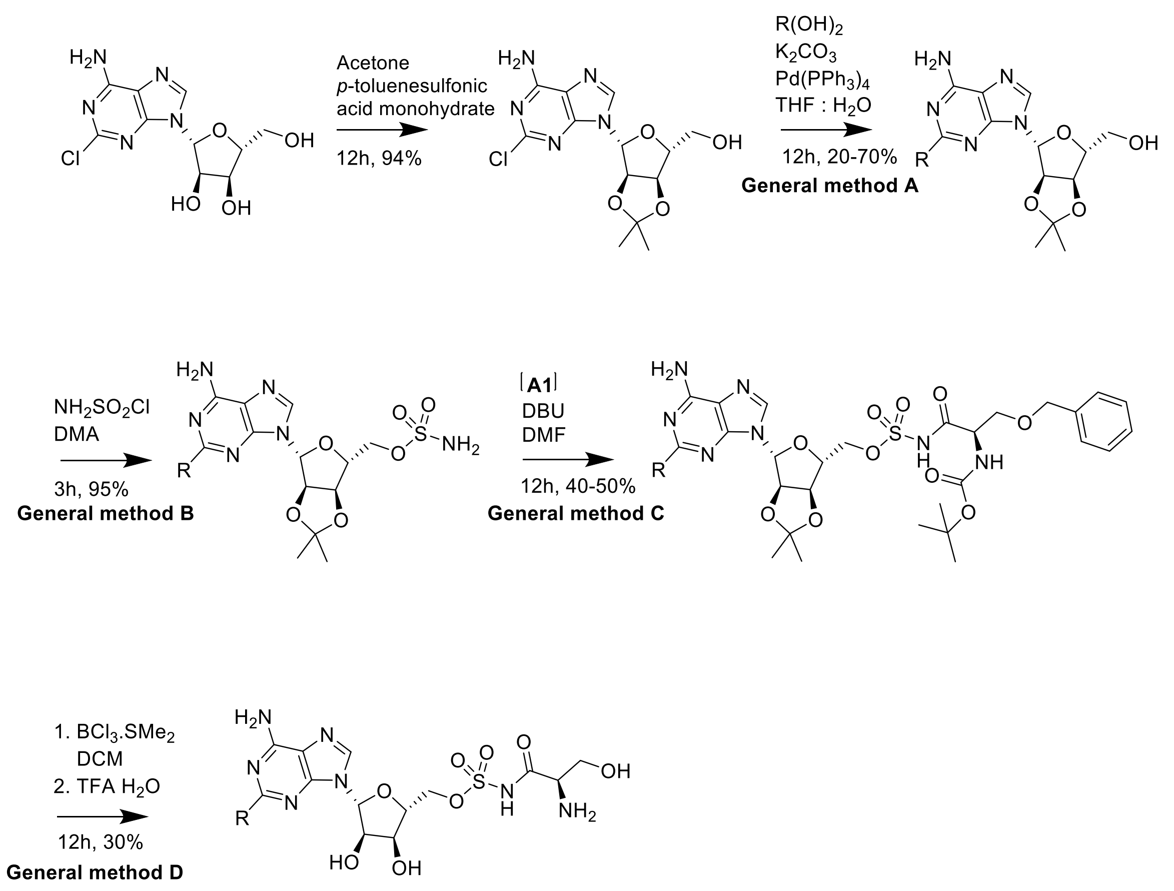


**2-8**

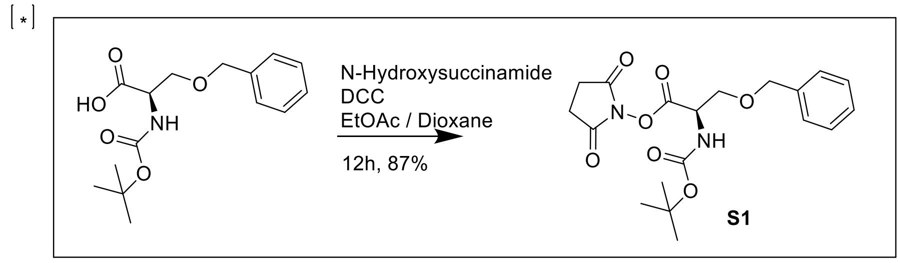


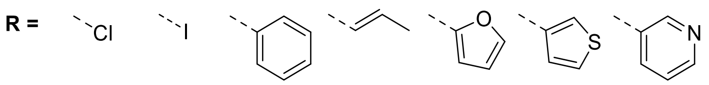


**2** **3** **4** **5** **6** **7** **8**

**General method A: Suzuki coupling reactions**

The boronic acid (4.0 equiv.) was added to a stirred solution of 2-chloroadenosine (1.0 equiv.), potassium carbonate (2.00 equiv.) and tetrakis (triphenylphosphine) palladium (0) (0.20 equiv.) in THF (8 mL) and water (4 mL). The reaction mixture was heated to reflux for 12 h. The reaction mixture was filtered through Celite® and concentrated *in vacuo*. The residue was diluted with water (20 mL) and extracted in EtOAc (3 × 20 mL). The combined organics were washed with water (3 × 10 mL) and brine (3 × 10 mL), dried ($MgSO_4$) and concentrated *in vacuo* to give an off-white/yellow solid, which was purified using flash column chromatography to afford the coupled products which were used without further purification.

**General method B: Introduction of sulfonyl group**

Generation of sulfanoyl chloride:

Chlorosulfonyl isocyanate (1 mL, 11.5 mmol, 1.0 equiv.) was cooled to 0 °C under an atmosphere of nitrogen. Formic acid (0.43 mL, 11.5 mmol, 1.0 equiv.) was added dropwise and the mixture stirred at room temperature overnight. Gas evolution was observed. The resulting colourless solid was dried *in vacuo*. The colourless solid was used without further purification / characterisation.

Sulfanoyl chloride (2.0 equiv.) in DMA (2 mL) was added dropwise to a stirred mixture of acetyl protected adenolate (1.0 equiv.) in DMA (3 mL) at 0 °C under an atmosphere of nitrogen. The mixture was then stirred at room temperature for 3 h. The reaction mixture was quenched with $Et_3N$ (1.5 mL) then MeOH (5 mL). The resulting solution was concentrated *in vacuo* before EtOAc (50 mL) was added. The mixture was extracted with 5% $NaHCO_3$ (3 × 15 mL), brine (3 × 10 mL), dried ($MgSO_4$) and concentrated *in vacuo* to afford the desired product as a colourless glassy solid.

**General method C: Amide coupling reaction**

Sulfonamide adenylate (1.0 equiv.) was dissolved in DMF (10 mL). After addition of DBU (1.1 equiv.), N-Boc-Ser(bzl)-OSu (1.1 equiv.) was added to the reaction mixture. After stirring for 16 h at room temperature, the mixture was concentrated *in vacuo* and the residue taken up in water (50 mL) and extracted with dichloromethane (50 mL). The organic layers were dried ($MgSO_4$) concentrated *in vacuo* and purified by flash chromatography (EtOAc) to afford the desired product as a colourless powder.

**General method D: Global deprotection**

Seryl-sulfonamide adenylate (1.0 equiv.) was dissolved in DCM (5 mL) under an atmosphere of nitrogen. To this was added $BCl_3.SMe_2$ (2M in DCM, 7.00 equiv.) and the reaction was stirred at room temperature for 8 h. The mixture was concentrated *in vacuo*. The residue was re-suspended in TFA:$H_2O$ (3:1, 4 mL) and the reaction was stirred overnight at room temperature. The mixture was concentrated *in vacuo*. The crude product was purified by preparative HPLC to afford the desired product as a colourless solid.

**Preparation of 2'3'-O-Isopropylidene-2-chloroadenosine**

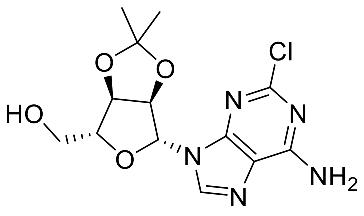


2-Chloroadenosine (0.20 g, 0.66 mmol, 1.0 equiv.) and p-toluenesulfonic acid mono hydrate (1.26 g, 6.6 mmol, 10 equiv.) were dissolved in acetone (100 mL) under an atmosphere of nitrogen. The reaction was stirred at room temperature overnight. While cooling in an ice bath a saturated $NaHCO_3$ solution (100 mL) was added to the reaction mixture until the pH of the solution was slightly basic. The acetone was removed *in vacuo*. The remaining aqueous solution was extracted with EtOAc (3 × 50 mL). The combined organics were dried ($MgSO_4$) and concentrated *in vacuo*. The desired product was isolated as a colourless solid (203 mg, 0.59 mmol, 90%); m.p.: 184.2-185.6 °C; $R_f$: 0.67 (9:1 Chloroform-MeOH); $\delta_H$ (500 MHz, DMSO-d6): 8.37 (1H, s, 6-HAr), 7.87 (2H, brs, $NH_2$), 6.07 (1H, d, *J* 2.8 2-HFuryl), 5.29 (1H, dd, *J* 6.2 and 2.8, 3-HFuryl), 5.09 (1H, app t, *J* 5.4, OH), 4.95 (1H, dd, *J* 6.2 and 2.8, 4-HFuryl), 4.22 (1H, dd, *J* 6.2 and 5.4, 5-HFuryl), 3.65-3.50 (2H, m, $CH_2$), 1.56 (3H, s, $CH_3$a), 1.34 (3H, s, $CH_3$b); $\delta_C$ (125 MHz, DMSO-d6): 158.1 (C4Ar), 153.0 (C2Ar), 149.9 (C8Ar), 139.9 (C6Ar), 128.2 (C5Ar), 119.0 (C9Ar), 113.1 (Acetyl C), 89.4 (C2Furyl), 86.7 (C5Furyl), 83.4 (C3Furyl), 81.2 (C4Furyl), 61.5 ($CH_2$), 37.0 ($CH_3$a), 25.2 ($CH_3$b); $\nu_{max}$/ $cm^{-1}$ (solid): 3473, 3299, 1761, 1651, 1381; HPLC: $T_r$= 2.26 (100% rel. area); *m/z* (ES): (Found: $[M+H]^+$, 342.0965. $C_{13}H_{16}ClN_5O_4$ requires *[M+H]*, 342.0964. $[\alpha]_D$ = -115.5° (c 0.1, MeOH).

**Preperation of [(3aR, 4R, 6R, 6aR)-6-(6-amino-2-chloro-9H-purin-9-yl)-2,2-dimethyl-tetrahydro-2H-furo[3,4-d][1,3]dioxol-4-yl]methyl sulfamate**

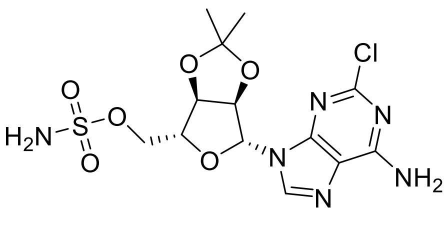


Preparation was *via* general method B using 2'3'-O-Isopropylidene-2-chloroadenosine (0.17 g, 0.49 mmol, 1.0 equiv.) to afford the desired product as a colourless glassy solid (0.19 g, 0.46 mmol, 94%) m.p.: 69.4-71.7 °C; $R_f$: 0.46 (9:1 Chloroform-MeOH); $\delta_H$ (500 MHz, DMSO-d6): 8.33 (1H, s, 6-HAr), 7.90 (2H, brs, $NH_2$), 7.60 (2H, brs, $SNH_2$), 6.18 (1H, d, *J* 2.5, 2-HFuryl), 5.36 (1H, dd, *J* 6.2 and 2.3, 3-HFuryl), 5.03 (1H, dd, *J* 6.2 and 3.4, 4-HFuryl), 4.43 (1H, dd, *J* 9.0 and 3.4, 5-HFuryl), 4.23 (2H, ddd, *J* 17.2, 9.0 and 3.4, $CH_2$), 1.57 (3H, s, $CH_3$a), 1.36 (3H, s, $CH_3$b); $\delta_C$ (125 MHz, DMSO-d6): 156.9 (C4Ar), 153.2 (C2Ar), 149.8 (C8Ar), 139.9 (C6Ar), 118.1 (C9Ar), 113.7 (C Acetyl), 88.8 (C2Furyl), 83.7 (C4Furyl), 83.4 (C3Furyl), 80.9 (C4Furyl), 68.1 ($CH_2$), 26.9 ($CH_3$a), 25.2 ($CH_3$b); $\nu_{max}$/

cm$^{-1}$ (solid): 3321, 3171, 1594, 1206; HPLC: $T_r$= 2.30 (95% rel. area); *m/z* (ES): (Found: $[M+H]^+$, 421.0695. $C_{13}H_{17}ClN_6O_6S$ requires *[M+H]*, 421.0692. $[\alpha]_D$ = -20.2° (c 0.1, MeOH).

**Preparation of N-Boc-Ser(bzl)-OSu (S1)**

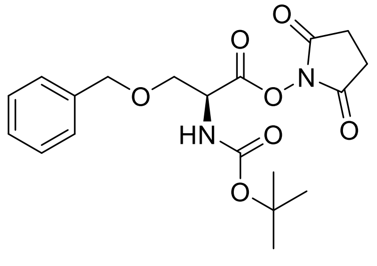


To a stirred solution of N-Boc-(Bzl)-Ser-OH (0.50 g, 1.69 mmol, 1.0 equiv.) in EtOAc/Dioxane (1:1, 10 mL) cooled to 0 °C were added N-hydroxysuccinimide (0.21 g, 1.78 mmol, 1.05 equiv.) and DCC (0.37 g, 1.78 mmol, 1.05 equiv.). The resulting mixture was stirred at room temperature overnight. The reaction mixture was filtered through Celite® and the filtrate concentrated *in vacuo*. The residue was re-suspended in EtOAc (35 mL), washed with 5% $NaHCO_3$ (3 × 5 mL), water (2 × 10 mL), and brine (10 mL). The organic phase was dried ($MgSO_4$) and concentrated in vacuo to afford the desired product as a colourless solid which was used without further purification. (0.35 g, 0.90 mmol, 53 %) m.p.:98.3-99.2 °C; $R_f$: 0.5 (9:1 DCM-MeOH); $\delta_H$ (500 MHz, DMSO-d6): 7.45-7.28 (5H, m, H-Bzl), 4.69 (1H, d, J 5.7, H-Chiral), 4.55 (2H, s, $CH_2$Bzl), 3.80 (2H, d, J 5.7, CH2), 2.82 (4H, brs, $CH_2$C=O), 1.41 (9H, s, BOC); $\delta_C$ (125 MHz, DMSO-d6): 169.8 (Succinimide C=O), 166.8 (C=O), 155.2 (BOC C=O), 137.8 (C1Ar), 128.2 (C3 and C5), 127.5 (C4), 127.5 (C2 and C6), 78.9 (C BOC), 72.3 ($CH_2$Bzl), 68.6 ($CH_2$C), 52.3 (Chiral C), 28.1 ($CH_3$ × 3), 25.4 ($CH_2$ × 2); $\nu_{max}$/ cm$^{-1}$ (solid): 3366, 2970, 1741, 1518, 1366; HPLC: $T_r$= 2.61 (83% rel. area); *m/z* (ES): (Found: $[M+H]^+$, 393.1663. $C_{19}H_{24}N_2O_7$ requires *[M+H]*, 393.1656. $[\alpha]_D$ = -7.6° (c 0.1, MeOH).

**Preparation of tert-butyl N-[(2S)-1-[((([(3aR, 4R, 6R, 6aR)-6-(6-amino-2-chloro-9H-purin-9-yl)-2,2-dimethyl-tetrahydro-2H-furo[3,4-d][1,3]dioxol-4-yl]methoxy)sulfonyl)amino]-3-(benzyloxy)-1-oxopropan-2-yl]carbamate**

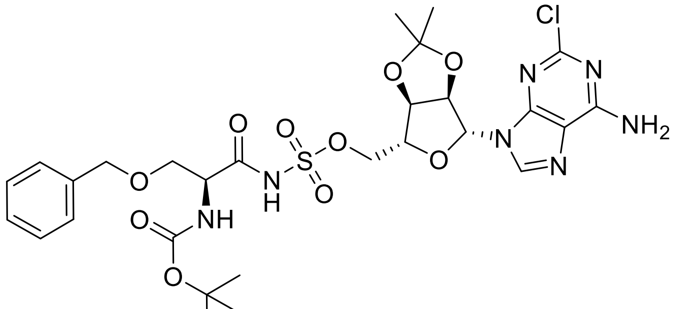


Preparation was *via* general method C using [(3aR, 4R, 6R, 6aR)-6-(6-amino-2-chloro-9H-purin-9-yl)-2,2-dimethyl-tetrahydro-2H-furo[3,4-d][1,3]dioxol-4-yl]methyl sulfamate (0.15 g, 0.36 mmol) to afford the desired product as a colourless glassy solid (0.19 g, 0.46 mmol, 94%) m.p.: 168.6-170.2 °C; $R_f$: 0.51 (9:1 DCM-MeOH); $\delta_H$ (500 MHz, DMSO-d6): 8.43 (1H, s, 6-HAr), 7.83 (2H, brs, NH2), 7.30-7.20 (5H, m, Bzl), 6.10 (1H, d, *J* 5.7, 2-HFuryl), 6.04 (1H, brs, NH), 5.25 (1H, d, *J* 5.7, 3-HFuryl), 4.94 (1H, d, *J* 5.7, 4-HFuryl), 4.47 (2H, s, $CH_2$ Bzl), 4.35 (1H, d, *J* 5.7, 5-HFuryl), 4.07-4.01 (3H, m, $CH_2$Chiral and Chiral H), 3.69-3.67 (2H, m, $CH_2$), 1.55 (3H, s, $CH_3$a), 1.29 (3H, s, $CH_3$b); $\delta_C$ (125 MHz, DMSO-d6): 171.6 (C=O), 157.8 (C4Ar), 155.4 (C=O BOC), 153.9 (C2Ar), 151.0

(C8Ar), 140.3 (C6Ar), 138.0 (C1Bzl), 128.7 (C3Bzl and C5Bzl), 128.3 (C2Bzl and C6Bzl), 127.9 (C4Bzl), 119.1 (C9Ar), 114.4 (C acetate), 90.6 (C2Furyl) 84.4 (C5Furyl), 83.3 (C3Furyl), 81.4 (C4Furyl), 79.5 (BOC C), 73.7 ($CH_2$ Bzl), 70.5 ($CH_2O$), 69.8 ($CH_2C$), 54.9 (Chiral C), 28.3 (CH3 × 3), 26.7 ($CH_3$a), 24.9 ($CH_3$b); $\nu_{max}$/ $cm^{-1}$ (solid):3330, 1691, 1637, 1304; HPLC: $T_r$= 2.59 (100% rel. area); *m/z* (ES): (Found: $[M+H]^+$, 698.2012. $C_{28}H_{36}ClN_7O_{10}S$ requires *[M+H]*, 698.2006. $[\alpha]_D$ = -50.4° (c 0.1, MeOH).

**Preparation of (2S)-2-amino-1-[([[(2R, 3S, 4R, 5R)-5-(6-amino-2-chloro-9H-purin-9-yl)-3,4-dihydroxyoxolan-2-yl]methoxy)sulfonyl)amino]-3-hydroxypropan-1-one (2)**

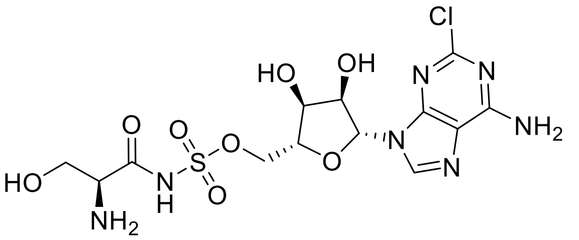

Preparation was via general method D using tert-butyl N-[(2S)-1-[((([(3aR, 4R, 6R, 6aR)-6-(6-amino-2-chloro-9H-purin-9-yl)-2,2-dimethyl-tetrahydro-2H-furo[3,4-d][1,3]dioxol-4-yl]methoxy)sulfonyl)amino]-3-(benzyloxy)-1-oxopropan-2-yl]carbamate (0.30 g, 0.43 mmol) to afford the desired product as a colourless powder (50.4 mg, 0.11 mmol, 25%) m.p.: 121.3 °C (Decomp); $R_f$: Baseline (9:1 DCM-MeOH); $\delta_H$ (500 MHz, DMSO-d6): 8.04 (1H, s, 6-HAr), 7.65 (2H, brs, $NH_2$), 5.97 (1H, d, *J* 8.2, 2-HFuryl), 4.60 (1H, apps, 3-HFuryl), 4.31 (1H, dd, *J* 10.1 and 5.9, $CH_2$*O), 4.19-4.15 (2H, m, $CH_2$*O and 4-HFuryl), 4.13 (1H, app s, 5-HFuryl), 3.81 (1H, dd, *J* 10.2 and 7.3, $CH_2$*chiral), 3.76 (1H, d, *J* 7.3, CHChiral), 3.65 (1H, dd, *J* 10.2 and 7.3, $CH_2$*chiral); $\delta_C$ (125 MHz, DMSO-d6): 172.1 (C=O), 157.8 (C4Ar), 154.1 (C2Ar), 151.0 (C9Ar), 140.1 (C6Ar), 119.1 (C10Ar), 89.4 (C2Furyl), 84.4 (C5Furyl), 73.9 (C3Furyl), 71.5 (C4Furyl), 70.3 ($CH_2O$), 64.4 ($CH_2$Chiral), 56.9 (CHChiral); $\nu_{max}$/ $cm^{-1}$ (solid): 3321, 3125, 2784, 1673, 1592, 1358; *m/z* (ES): No mass ion found $[\alpha]_D$ = 28.8° (c 0.1, MeOH).

**Preparation of 2'3'-O-Isopropylidene-2-iodoadenosine**

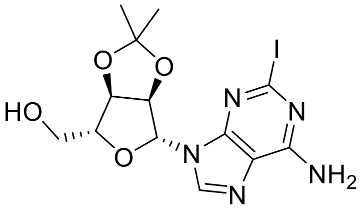

2-iodoadenosine (1.00 g, 2.54 mmol, 1.0 equiv.) and p-toluenesulfonic acid mono hydrate (4.83 g, 25.4 mmol, 10 equiv.) were dissolved in acetone (200 mL) under an atmosphere of nitrogen. The reaction was stirred at room temperature overnight. While cooling in an ice bath a saturated $NaHCO_3$ solution (200 mL) was added to the reaction mixture until the pH of the solution was slightly basic. The acetone was removed *in vacuo*. The remaining aqueous solution was extracted with EtOAc (3 × 100 mL). The combined organics were dried ($MgSO_4$) and concentrated *in vacuo*. The desired product was isolated as a colourless solid (0.94 g, 2.17 mmol, 86%); m.p.: 182.4-184.1 °C; $R_f$: 0.69 (9:1 Chloroform-MeOH); $\delta_H$ (400 MHz, DMSO-d6): 8.34 (1H, s, 6-HAr), 7.83 (2H, brs, $NH_2$), 6.06 (1H, d, *J* 4.0, 2-HFuryl), 5.30 (1H, dd, J 6.2 and 4.0, 3-HFuryl), 5.09 (1H, app t, *J* 5.2, OH), 4.96 (1H, dd, *J* 6.2 and 3.9, 4-HFuryl), 4.20 (1H, d, *J* 3.9, 5-

HFuryl), 3.52 (2H, app s, $CH_2$), 1.56 (3H, s, $CH_3$a), 1.37 (3H, s, $CH_3$b); $\delta_C$ (100 MHz, DMSO-d6): 158.2 (C4Ar), 156.4 (C8Ar), 149.8 (C6Ar), 121.4 (C8Ar), 119.3 (C2Ar) 113.6 (Acetyl C), 89.6 (C2Furyl), 87.3 (C5Furyl), 84.0 (C3Furyl), 81.7 (C4Furyl), 62.0 ($CH_2$), 27.5 ($CH_3$a), 25.7 ($CH_3$b); $\nu_{max}$/ $cm^{-1}$ (solid): 3473, 3299, 1761, 1651, 1381; HPLC: $T_r$= 2.30 (100% rel. area); *m/z* (ES): (Found: $[M+Na]^+$, 455.8. $C_{13}H_{16}IN_5O_4$ requires *[M+Na]*, 455.8. $[\alpha]_D$ = -102.0° (c 0.1, MeOH).

**Preperation of [(3aR, 4R, 6R, 6aR)-6-(6-amino-2-iodo-9H-purin-9-yl)-2,2-dimethyl-tetrahydro-2H-furo[3,4-d][1,3]dioxol-4-yl]methyl sulfamate**

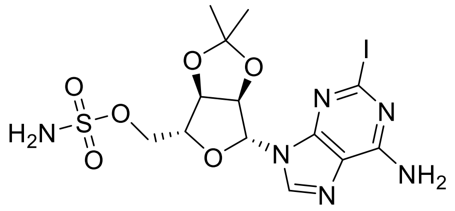


Preparation was *via* general method B using 2'3'-O-Isopropylidene-2-iodoadenosine (0.20 g, 0.46 mmol) to afford the desired product as a colourless glassy solid (0.20 g, 0.39 mmol, 85%) m.p.: 70.2-71.7 °C; $R_f$: 0.48 (9:1 Chloroform-MeOH); $\delta_H$ (500 MHz, DMSO-d6): 8.24 (1H, s, 6-HAr), 7.80 (2H, brs, $NH_2$), 7.59 (2H, brs, $SNH_2$), 6.18 (1H, d, *J* 4.0, 2-HFuryl), 5.33 (1H, dd, *J* 6.1 and 4.0, 3-HFuryl), 5.02 (1H, dd, *J* 6.1 and 3.8), 4.43 (1H, app d, *J* 3.8, 5-HFuryl), 4.28-4.12 (2H, m, $CH_2$), 1.63 (3H, s, $CH_3$a), 1.38 (3H, s, $CH_3$b); $\delta_C$ (100 MHz, DMSO-d6): 156.5 (C4Ar), 156.4 (C8Ar), 149.7 (C6Ar), 121.5 (C9Ar), 119.4 (C2Ar), 114.2 (Acetyl C), 89.1 (C2Furyl), 84.3 (C5Furyl), 84.1 (C3Furyl), 81.4 (C4Furyl), 68.5 ($CH_2$), 27.4 ($CH_3$a), 25.7 ($CH_3$b); $\nu_{max}$/ $cm^{-1}$ (solid): 3321, 3171, 1594, 1206; HPLC: $T_r$= 2.35 (100% rel. area); *m/z* (ES): (Found: $[M+Na]^+$, 442.9. $C_{13}H_{17}IN_6O_6S$ requires *[M+Na]*, 442.9. $[\alpha]_D$ = -21.9 ° (c 0.1, MeOH).

**Preparation of tert-butyl N-[(2S)-1-[(([(3aR, 4R, 6R, 6aR)-6-(6-amino-2-iodo-9H-purin-9-yl)-2,2-dimethyl-tetrahydro-2H-furo[3,4-d][1,3]dioxol-4-yl]methoxy)sulfonyl)amino]-3-(benzyloxy)-1-oxopropan-2-yl]carbamate**

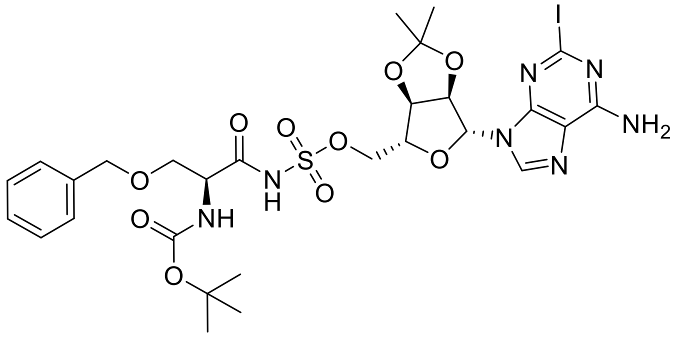


Preparation was *via* general method C using [(3aR, 4R, 6R, 6aR)-6-(6-amino-2-iodo-9H-purin-9-yl)-2,2-dimethyl-tetrahydro-2H-furo[3,4-d][1,3]dioxol-4-yl]methyl sulfamate (0.20 g, 0.39 mmol) to afford the desired product as a colourless glassy solid (40.4 mg, 0.05 mmol, 13%) m.p.: 166.2-168.9°C; $R_f$: 0.54 (DCM-MeOH); $\delta_H$ (400 MHz, DMSO-d6): 8.38 (1H, s, 6-HAr), 7.75 (2H, brs, $NH_2$), 7.30-7.20 (5H, m, Bzl), 6.07 (1H, app s, 2-HFuryl), 5.24 (1H, app s, 3-HFuryl), 4.92 (1H, app s, 4-HFuryl), 4.42 (2H, app s, $CH_2$ Bzl), 4.35 (1H, app s, 5-HFuryl), 4.08-4.01 (3H, m, $CH_2$ Chiral and Chiral H) 3.70-3.62 (2H, m, $CH_2$), 1.54 (3H, s, $CH_3$a), 1.38 (9H, s, CH3 × 3), 1.20 (3H, s, $CH_3$b); $\delta_C$ (100 MHz, DMSO-d6): 174.5 (C=O), 157.3 (C4Ar), 155.5 (C=O BOC), 152.3 (C8Ar), 139.0 (C6Ar),

137.7 (C1Bzl), 128.7 (C5Bzl and C3Bzl), 128.1 (C4Bzl), 127.8 (C2’Ar and C6’Ar), 118.5 (C9Ar), 116.4 (C2Ar), 114.4 (Acetyl C), 88.3 (C2Furyl), 82.8 C5Furyl), 81.5 (C3Furyl), 79.6 (BOC C), 79.0 (C4Furyl), 73.5 ($CH_2$ Bzl); 69.9 ($CH_2O$), 67.4 ($CH_2C$), 56.4 (Chiral C), 28.3 ($CH_3$ × 3), 26.4 ($CH_3$a), 24.9 ($CH_3$b); $\nu_{max}$/ $cm^{-1}$ (solid): 3330, 1691, 1637, 1304; HPLC: $T_r$= 2.67 (100% rel. area); *m/z* (ES): (Found: $[M+Na]^+$, 812.1. $C_{28}H_{36}IN_7O_{10}S$ requires *[M+Na]*, 812.1. $[\alpha]_D$ = -52.2° (c 0.1, MeOH).

**Preparation of (2S)-2-amino-1-[([[(2R, 3S, 4R, 5R)-5-(6-amino-2-iodo-9H-purin-9-yl)-3,4-dihydroxyoxolan-2-yl]methoxy)sulfonyl)amino]-3-hydroxypropan-1-one (3)**

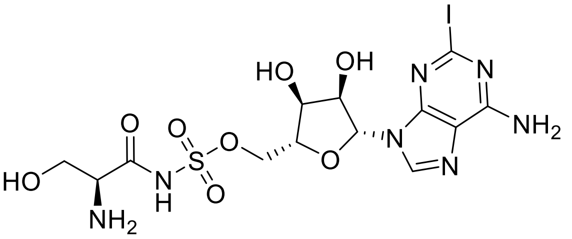

Preparation was via general method D using tert-butyl N-[(2S)-1-[(([(3aR, 4R, 6R, 6aR)-6-(6-amino-2-iodo-9H-purin-9-yl)-2,2-dimethyl-tetrahydro-2H-furo[3,4-d][1,3]dioxol-4-yl]methoxy)sulfonyl)amino]-3-(benzyloxy)-1-oxopropan-2-yl]carbamate (50 mg, 0.06 mmol) to afford the desired product as a colourless powder (4.0 mg, 0.07 mmol, 12%) m.p.: 135.2 °C (Decomp); $R_f$: Baseline (9:1 DCM-MeOH); $\delta_H$ (500 MHz, DMSO-d6): 8.36 (1H, s, 6-HAr), 7.68 (2H, brs, $NH_2$), 5.88 (1H, d, J 5.4, 2-HFuryl), 4.60 (1H, apps, 3-HFuryl), 4.31 (1H, dd, *J* 10.1 and 5.9, $CH_2$*O), 4.19-4.15 (2H, m, $CH_2$*O and 4-HFuryl), 4.13 (1H, app s, 5-HFuryl), 3.81 (1H, dd, *J* 10.2 and 7.3, $CH_2$*chiral), 3.76 (1H, d, *J* 7.3, CH Chiral), 3.65 (1H, dd, *J* 10.2 and 7.3, $CH_2$*chiral); $\nu_{max}$/ $cm^{-1}$ (solid): 3321, 3125, 2784, 1673, 1592, 1358; *m/z* (ES): (Found: $[M-H]^-$, 558.0. $C_{13}H_{18}IN_7O_8S$ requires *[M-H]*, 558.0. $[\alpha]_D$ = 28.4° (c 0.1, MeOH).

**Preparation of 2’3’-O-Isopropylidene-2-phenyladenosine**

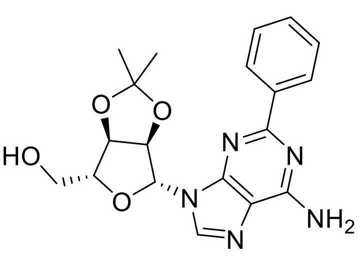

Preparation was via general method A using 2’3’-O-Isopropylidene-2-chloroadenosine (0.50 g, 1.46 mmol) and phenyl boronic acid (0.71 g, 5.85 mmol) to afford the desired product as a colourless powder (0.43 g, 1.12 mmol, 76%); m.p.: 166.2-168.1 °C; $R_f$: 0.58 (9:1 Chloroform-MeOH); $\delta_H$ (500 MHz, DMSO-d6): 8.36 (3H, app s, 6-HAr, 2’-HAr and 6’-HAr), 7.46 (3H, app s, 4’-HAr, 3’-HAr and 5’-HAr), 7.39 (2H, brs, $NH_2$), 6.26 (1H, app s, 2-HFuryl), 5.52 (1H, app s, 3-HFuryl), 5.10 (1H, app s, 4-HFuryl), 5.04 (1H, app s, OH), 4.22 (1H, app s, 4-HFuryl), 3.69-3.51 (2H, m, $CH_2$), 1.59 (3H, s, $CH_3$a), 1.37 (3H, s, $CH_3$b); $\delta_C$ (125 MHz, DMSO-d6): 158.5 (C2Ar), 156.4 (C4Ar), 150.4 (C8Ar), 140.8 (C6Ar), 138.8 (C1’Ar), 134.4 (C4’Ar), 128.7 (C3’Ar and C5’Ar), 128.2 (C2’ Ar and C6’Ar), 118.7 (C9Ar), 113.6 (Acetyl C), 89.5 (C2Furyl), 87.2 (C5Furyl), 83.8 (C3Furyl), 81.9 (C4Furyl), 62.0 ($CH_2$), 27.6 ($CH_3$a), 25.7 ($CH_3$b); $\nu_{max}$/ $cm^{-1}$ (solid): 3314, 3157, 1655, 1596, 1372; HPLC: $T_r$= 2.24 (100% rel. area); *m/z* (ES): (Found: $[M+H]^+$, 384.1676. $C_{19}H_{21}N_5O_4$ requires *[M+H]*, 384.1666. $[\alpha]_D$ = -1.4° (c 0.1, MeOH).

**Preperation of [(3aR, 4R, 6R, 6aR)-6-(6-amino-2-phenyl-9H-purin-9-yl)-2,2-dimethyl-tetrahydro-2H-furo[3,4-d][1,3]dioxol-4-yl]methyl sulfamate**

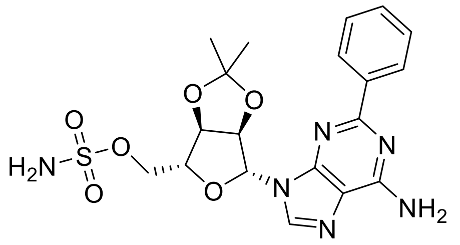

Preparation was *via* general method B using 2'3'-O-Isopropylidene-2-phenyladenosine (0.40 g, 1.04 mmol) to afford the desired product as a colourless glassy solid (0.23 g, 0.50 mmol, 48%) m.p.: 89.9-91.2 °C; $R_f$: 0.50 (9:1 Chloroform-MeOH); $\delta_H$ (500 MHz, DMSO-d6): 8.38-8.33 (3H, m, 6-HAr, 2'-HAr and 6'-HAr), 7.60 2H, brs, $SNH_2$), 7.54-7.45 (3H, m, 3'-HAr, 4'-HAr and 5'-HAr), 7.42 2H, brs, $NH_2$), 6.34 (1H, app s, 2-HFuryl), 5.36 1H, d, J 6.3, 3-HFuryl), 5.20 (1H, app s, 4-HFuryl), 4.46 (1H, app s, 5-HFuryl), 4.25-4.15 (2H, m, $CH_2$), 1.60 (3H, s, $CH_3$a), 1.38 (3H, s, $CH_3$b); $\delta_C$ (125 MHz, DMSO-d6): 158.6 (C2Ar), 156.4 (C4Ar), 150.3 (C8Ar), 140.8 (C6Ar), 138.7 (C1'Ar), 130.2 (C4'Ar), 129.2 (C3'Ar and C5'Ar), 128.7 (C6'Ar and C2'Ar), 89.3 (C2Furyl), 84.1 (C5Furyl), 83.8 (C3Furyl), 81.6 (C4Furyl), 68.6 ($CH_2$), 27.5 ($CH_3$a), 25.7 ($CH_3$b); $\nu_{max}$/ $cm^{-1}$ (solid): 3354, 2936, 1628, 1376; HPLC: $T_r$= 2.27 (100% rel. area); *m/z* (ES): (Found: $[M+H]^+$, 463.1401. $C_{19}H_{22}N_6O_6S$ requires *[M+H]*, 463.1394. $[\alpha]_D$ = 17.6° (c 0.1, MeOH).

**Preparation of tert-butyl N-[(2S)-1-[((([(3aR, 4R, 6R, 6aR)-6-(6-amino-2-phenyl-9H-purin-9-yl)-2,2-dimethyl-tetrahydro-2H-furo[3,4-d][1,3]dioxol-4-yl]methoxy)sulfonyl)amino]-3-(benzyloxy)-1-oxopropan-2-yl]carbamate**

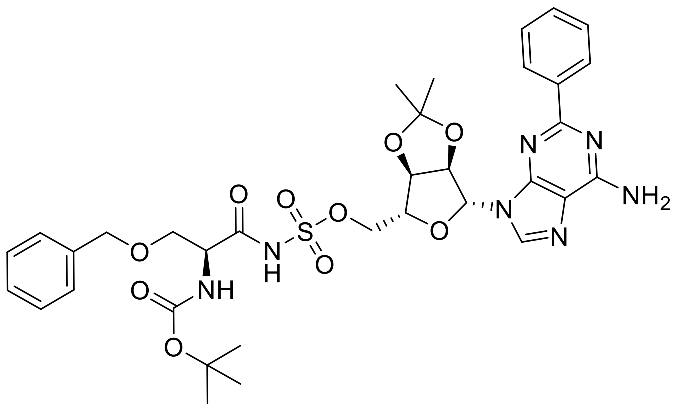

Preparation was *via* general method C using [(3aR, 4R, 6R, 6aR)-6-(6-amino-2-phenyl-9H-purin-9-yl)-2,2-dimethyl-tetrahydro-2H-furo[3,4-d][1,3]dioxol-4-yl]methyl sulfamate (0.20 g, 0.43 mmol) to afford the desired product as a colourless glassy solid (0.21 g, 0.29 mmol, 67%) m.p.: 155.8-157.7 °C; $R_f$: 0.64 (9:1 DCM-MeOH); $\delta_H$ (500 MHz, DMSO-d6): 8.42(1H, s, 6-HAr), 8.37 (2H, d, J 6.9 6'-HAr and 2'-HAr), 7.57-7.41 (3H, m, 5'-HAr, 4'-HAr and 3'-HAr), 7.38 (2H, brs, $NH_2$), 7.26-7.24 (5H, m, 2-HBzl, 3-HBzl, 4-HBzl, 5-HBzl and 6-HBzl), 6.27 (1H, d, J 3.0, 2-HFuryl), 6.06 (1H, d J 8.1, NH), 5.46 (1H, d, J 3.0, 3-HFuryl), 5.07 (1H, app s, 4-HFuryl), 4.45-4.41 (3H, m, BzlCH2, 5-HFuryl), 4.05 (2H, d, J 4.9, CH2O), 3.96-3.92 (1H, m, Chiral H), 3.67-3.58 (2H, m, Chiral $CH_2$), 1.59 (3H, s, $CH_3$a), 1.39 (9H, s, $CH_3 \times 3$), 1.35 (3H, s, $CH_3$b); $\delta_C$ (125 MHz, DMSO-d6): 173.5 (C=ONH), 158.7 (C2Ar), 156.8 (C4Ar), 156.4 (C=O BOC), 150.6 (C8Ar), 140.0 (C6Ar), 138.5 (C1'Ar and C1Bzl), 130.4 (C4'Ar), 129.6-127.1 (C2Bzl, C3Bzl, C4Bzl, C5Bzl, C6Bzl,

C2'Ar, C3'Ar, C5'Ar and C6'Ar), 118.0 (C9Ar), 112.5 (Acetyl C), 88.8 (C2Furyl), 83.5 (C5Furyl), 83.4 (C3Furyl), 81.6 (C4Furyl), 79.5 (BOC C), 71.8 ($CH_2$ Bzl), 71.1 (Chiral $CH_2$), 67.1 ($CH_2$), 57.1 (Chiral C), 28.2 ($CH_3$ × 3), 27.1 ($CH_3$a), 25.2 ($CH_3$b); $\nu_{max}$/ $cm^{-1}$ (solid): 3358, 3193, 2980, 1629, 1575, 1438; HPLC: $T_r$= 2.54 (100% rel. area); *m/z* (ES): (Found: $[M+H]^+$, 740.2718. $C_{34}H_{41}N_7O_{10}S$ requires *[M+H]*, 740.2708. $[\alpha]_D$ = 17.9° (c 0.1, MeOH).

**Preparation of (2S)-2-amino-1-[([[(2R, 3S, 4R, 5R)-5-(6-amino-2-phenyl-9H-purin-9-yl)-3,4-dihydroxyoxolan-2-yl]methoxy)sulfonyl)amino]-3-hydroxypropan-1-one (4)**

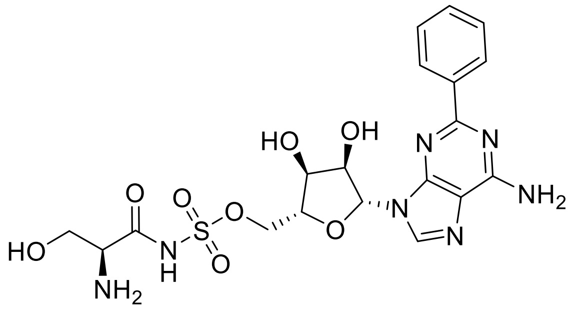

Preparation was via general method D using tert-butyl N-[(2S)-1-[((([(3aR, 4R, 6R, 6aR)-6-(6-amino-2-phenyl-9H-purin-9-yl)-2,2-dimethyl-tetrahydro-2H-furo[3,4-d][1,3]dioxol-4-yl]methoxy)sulfonyl)amino]-3-(benzyloxy)-1-oxopropan-2-yl]carbamate (0.20 g, 0.27 mmol) to afford the desired product as a colourless powder (54.0 mg, 0.11 mmol, 39%) m.p.: 143.2 (Decomp); *R*f: Baseline (9:1 DCM-MeOH); $\delta_H$ (500 MHz, DMSO-d6): 8.47 (1H, s, 6-HAr), 8.43 (2H, dd, J 8.0 and 1.6, 6'-HAr and 2'-HAr), 8.20 (1H, s, NH), 7.91 (2H, brs, NH2), 7.58-7.48 (3H, m, 3'-HAr, 4'-HAr and 5'-HAr), 6.09 (1H, d, J 5.9, 2-HFuryl), 4.77-4.70 (1H, m, 3-HFuryl), 4.34-4.28 (2H, m, 4-HFuryl and CH2*O), 4.22-4.16 2H, m, 5-HFuryl and CH2*O), 3.87 (1H, dd, J 11.2 and 3.7, CH2*Chiral) 3.69 (1H, dd, J 11.2 and 7.3, CH2*Chiral), 3.60-3.56 (1H, m, ChiralH); $\delta_C$ (125 MHz, DMSO-d6): 172.0 (C=O), 163.5 (C2Ar), 155.9 (C4Ar), 150.8 (C8Ar), 140.5 (C6Ar), 137.8 (C1'Ar), 130.1 (C4'Ar), 128.7 (C5'Ar and C3'Ar), 128.2 (C6'Ar and C2'Ar), 19.1 (C9Ar), 87.1 (C2Furyl), 82.8 (C5Furyl), 73.8 (C3Furyl), 71.3 (C4Furyl), 68.5 ($CH_2O$), 61.0 ($CH_2$Chiral), 57.6 (Chiral H); $\nu_{max}$/ $cm^{-1}$ (solid): 3367, 3070, 1673, 1598; HPLC: $T_r$= 1.89 (100% rel. area); *m/z* (ES): (Found: $[M+H]^+$, 510.1405. $C_{19}H_{23}N_7O_8S$ requires *[M+H]*, 510.1402). $[\alpha]_D$ = 25.7° (c 0.1, MeOH).

**Preparation of 2'3'-O-Isopropylidene-2-propenyladenosine**

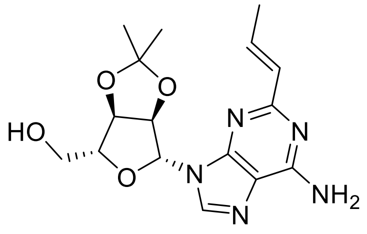

Preparation was via general method A using 2'3'-O-Isopropylidene-2-chloroadenosine (0.50 g, 1.46 mmol) and transpropenyl boronic acid (0.50 g, 5.85 mmol) to afford the desired product as a colourless powder (0.17 g, 0.50 mmol, 34%); m.p.: 81.6-83.4 °C; *R*f: 0.58 (9:1 Chloroform-MeOH); $\delta_H$ (500 MHz, DMSO-d6): 8.29 (1H, s, 6-HAr), 7.22 (2H, brs, $NH_2$), 6.92 (1H, dd, J 15.2 and 7.2, 2-Hpropyl), 6.33 (1H, d, J 15.2, 1-Hpropyl), 6.13 (1H, app s, 2-HFuryl), 5.34 (1H, app s, 3-HFuryl), 5.26 (1H, t, J 5.4, OH), 5.02 (1H, app s, 4-HFuryl), 4.22 (1H, app s, 5-HFuryl), 3.62-3.49 (2H, m, $CH_2$), 1.90 (3H, d, J 7.2, $CH_3$propyl), 1.56 (3H, s, $CH_3$a),

1.35 (3H, s, $CH_3$b); $\delta_C$ (125 MHz, DMSO-d6): 158.8 (C2Ar), 156.2 (C4Ar), 149.9 (C8Ar), 140.3 (C6Ar), 134.0 (C2Pro), 131.9 (C1Pro), 89.9 (C2Furyl), 86.9 (C5Furyl), 83.6 (C3Furyl), 81.9 (C4Furyl), 62.2 ($CH_2$), 27.6 ($CH_3$a), 25.7 ($CH_3$b), 18.3 ($CH_3$Pro); $\nu_{max}$/ $cm^{-1}$ (solid): 3451, 3310, 3162, 1657, 1575, 1373; HPLC: $T_r$= 2.11 (72% rel. area); *m/z* (ES): (Found: $[M+H]^+$, 348.3. $C_{16}H_{21}N_5O_4$ requires *[M+H]*, 348.5. $[\alpha]_D$ = -50.2° (c 0.1, MeOH).

**Preperation of [(3aR, 4R, 6R, 6aR)-6-(6-amino-2-propenyl-9H-purin-9-yl)-2,2-dimethyl-tetrahydro-2H-furo[3,4-d][1,3]dioxol-4-yl]methyl sulfamate**

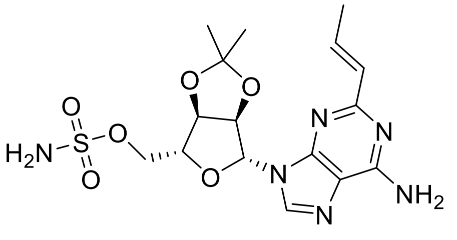


Preparation was *via* general method B using 2'3'-O-Isopropylidene-2-propenyladenosine (0.15 g, 0.43 mmol) to afford the desired product as a pale yellow oil (0.14 g, 0.33 mmol, 77%) $R_f$: 0.47 (9:1 Chloroform-MeOH); $\delta_H$ (500 MHz, DMSO-d6): 8.25 (1H, s, 6-HAr), 7.61 2H, brs, $SNH_2$), 7.23 (2H, brs, $NH_2$), 6.90 (1H, dd, J 13.4 and 6.6, 2-HPropyl), 6.36 (1H, d, J 13.4, 1-HPropyl), 6.24 (1H, app s, 2-HFuryl), 5.42 (1H, app s, 3-HFuryl), 5.15 (1H, app s, 4-HFuryl), 4.40 (1H, app s, 5-HFuryl), 4.35-4.25 (2H, m, $CH_2$), 1.90 (3H, d, J 6.6, $CH_3$ Propyl), 1.57 (3H, s, $CH_3$a), 1.36 (3H, s, $CH_3$b); $\delta_C$ (125 MHz, DMSO-d6): 171.0 (C2Ar), 156.1 (C4Ar), 150.9 (C8Ar), 140.0 (C6ar), 125.5 (C2Pro), 119.4 (C9Ar), 117.7 (C1Pro), 112.4 (Acetyl C), 88.2 (C2Furyl), 83.4 (C5Furyl), 83.1 (C3Furyl), 81.4 (C4Furyl), 70.5 ($CH_2$), 26.7 ($CH_3$a), 25.0 ($CH_3$b), 18.8 ($CH_3$Pro); $\nu_{max}$/ $cm^{-1}$ (solid): 3326, 3182, 2987, 2938, 1622, 1585, 1374; HPLC: $T_r$= 2.14 (67% rel. area); *m/z* (ES): (Found: $[M+H]^+$, 427.1400. $C_{16}H_{22}N_6O_6S$ requires *[M+H]*, 427.1394. $[\alpha]_D$ = 16.5° (c 0.1, MeOH).

**Preparation of tert-butyl N-[(2S)-1-[(([(3aR, 4R, 6R, 6aR)-6-(6-amino-2-propenyl-9H-purin-9-yl)-2,2-dimethyl-tetrahydro-2H-furo[3,4-d][1,3]dioxol-4-yl]methoxy)sulfonyl)amino]-3-(benzyloxy)-1-oxopropan-2-yl]carbamate**

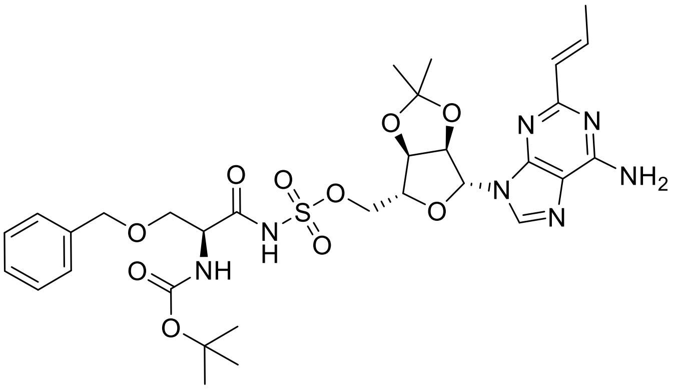


Preparation was *via* general method C using [(3aR, 4R, 6R, 6aR)-6-(6-amino-2-propenyl-9H-purin-9-yl)-2,2-dimethyl-tetrahydro-2H-furo[3,4-d][1,3]dioxol-4-yl]methyl sulfamate (0.14 g, 0.33 mmol) to afford the desired product as a colourless glassy solid (64.9 mg, 0.09 mmol, 28%) m.p.: 142.9-144.3 °C; $R_f$: 0.47 (9:1 DCM-MeOH); $\delta_H$ (500 MHz, DMSO-d6): 8.33 (1H, s, 6-HAr), 7.30-7.26 (5H, m, 2'-HAr, 3'-HAr, 4'-HAr, 5'-HAr and 6'-HAr), 7.18 (2H, brs, $NH_2$), 6.92 (1H, dd, J 15.3 and 7.0, 2-HPro), 6.35 (1H, d, J 15.3, 1-HPro), 6.15 (1H, d, J 2.8, 2-HFuryl), 6.06 (1H, d, J 7.7 NH), 5.33 (1H, dd, J 6.3 and 2.8, 3-HFuryl),

5.04 (1H, app t, J 6.3, 4-HFuryl), 4.42 (2H, d, J 9.9, $CH_2$ Bzl), 4.36 (1H, d J 2.4, 5-HFuryl), 4.15-3.95 (3H, m, $CH_2O$ and ChiralH), 3.73-3.54 (2H, m, $CH_2$ Chiral), 1.89 (3H, dd, J 7.0 and 1.5, Me), 1.56 (3H, s, $CH_3a$), 1.39 (9H, s, $CH_3 \times 3$), 1.33 (3H, s, $CH_3b$); $\delta_C$ (125 MHz, DMSO-d6): 172.5 (C2Ar), 170.1 (C=ONH), 156.1 (C4Ar), 155.4 (C=OBOC), 150.8 (C8Ar), 140.3 (C6Ar), 137.8 (C1'Ar), 128.7-127.5 (Aromatics), 125.2 (C2pro), 119.1 (C9Ar), 117.7 (C1Pro), 114.3 (Acetyl C), 88.9 (C2Furyl), 83.7 (C5Furyl), 83.4 (C3Furyl), 81.7 (C4Furyl), 79.5 (BOC C), 71.8 ($CH_2$ Bzl), 71.1 ($CH_2O$), 67.1 ($CH_2$ Chiral), 56.5 (Chiral C), 28.2 ($CH_3 \times 3$), 27.1 ($CH_3a$), 25.2 ($CH_3b$), 17.8 ($CH_3Pro$); $\nu_{max}$/ $cm^{-1}$ (solid): 3359, 2968, 1627, 1579, 1345; HPLC: $T_r$= 2.45 (78% rel. area); *m/z* (ES): (Found: $[M+H]^+$, 704.2719. $C_{31}H_{41}N_7O_{10}S$ requires *[M+H]*, 704.2708. $[\alpha]_D$ = -8.6° (c 0.1, MeOH).

**Preparation of (2S)-2-amino-1-[([[(2R, 3S, 4R, 5R)-5-(6-amino-2-propenyl-9H-purin-9-yl)-3,4-dihydroxyoxolan-2-yl]methoxy)sulfonyl)amino]-3-hydroxypropan-1-one (5)**

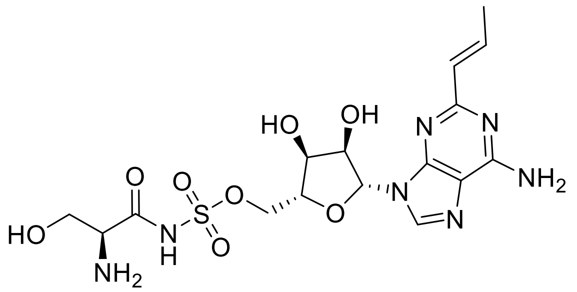


Preparation was via general method D using tert-butyl N-[(2S)-1-[(([(3aR, 4R, 6R, 6aR)-6-(6-amino-2-propenyl-9H-purin-9-yl)-2,2-dimethyl-tetrahydro-2H-furo[3,4-d][1,3]dioxol-4-yl]methoxy)sulfonyl)amino]-3-(benzyloxy)-1-oxopropan-2-yl]carbamate (65 mg, 0.09 mmol) to afford the desired product as a colourless powder (22.5 mg, 0.05 mmol, 53%) m.p.: 115.3 (Decomp); $R_f$: Baseline (9:1 DCM-MeOH); $\delta_H$ (500 MHz, DMSO-d6): 8.50 (1H, s, 6-HAr), 8.15 (1H, s, NH), 7.88 (2H, brs, $NH_2$), 7.09 (1H, dd, J 15.4 and 6.7, 2-HPropyl), 6.41 (1H, d, J 15.4, 1-HPropyl), 5.95 (1H, d, J 5.9, 2-HFuryl), 4.63 (1H, app t, J 5.9, 3-HFuryl), 4.28-4.09 (4H, m, 4-HFuryl, 5-HFuryl and $CH_2O$), 3.83 (1H, dd, J 11.1 and 3.6, $CH_2$* Chiral), 3.65 (1H, dd, 11.1 and 7.0, $CH_2$* Chiral), 3.56-3.50 (1H, m, Chiral H), 2.88 (2H, app d, J 5.3, $NH_2$), 1.98 (3H, d, J 6.7, $CH_3$); $\delta_C$ (125 MHz, DMSO-d6): 172.2 (C=O), 163.5 (C2Ar), 153.8 (C4Ar), 150.9 (C8Ar), 145.9 (C6Ar), 122.3 (C2Pro), 119.4 (C9Ar), 117.9 (C1Pro), 93.7 (C2furyl), 83.4 (C5Furyl), 75.4 (C3Furyl), 70.5 (C4Furyl), 70.2 ($CH_2O$), 57.3 ($CH_2$ Chiral), 54.8 (Chiral H), 19.1 ($CH_3$); $\nu_{max}$/ $cm^{-1}$ (solid): 3106, 2942, 1668, 1595, 1182; HPLC: $T_r$= 0.81 (100% rel. area); *m/z* (ES): (Found: $[M+H]^+$, 474.1406. $C_{16}H_{23}N_7O_8S$ requires *[M+H]*, 474.1402). $[\alpha]_D$ = 27.0° (c 0.1, MeOH).

**Preparation of 2'3'-O-Isopropylidene-2-(furan-2-yl)adenosine**

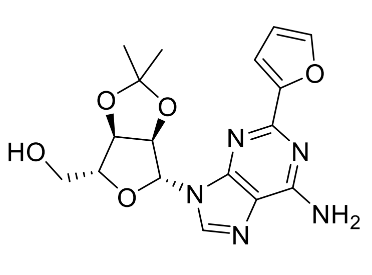


Preparation was via general method A using 2'3'-O-Isopropylidene-2-chloroadenosine (0.50 g, 1.46 mmol) and 2-furyl boronic acid (0.66 g, 5.85 mmol) to afford the desired product as a colourless powder (0.25 g, 0.67 mmol, 46%); m.p.: 96.3-98.2 °C; $R_f$: 0.59 (9:1 Chloroform-MeOH); $\delta_H$ (500 MHz, DMSO-d6): 8.35 (1H, s, 6-HAr),

7.83 (1H, 5'-HAr), 7.44 (2H, brs, NH2), 7.13 (1H, s, 3'-HAr), 6.65 (1H, s, 4'-HAr), 6.20 (1H, app s, 2-HFuryl), 5.40 (1H, app s, 3-HFuryl), 5.10 (1H, app s, 4-HFuryl), 5.05 (1H, app s, OH), 4.22 (1H, app s, 5-HFuryl), 3.67-3.52 (2H, m, $CH_2$), 1.55 (3H, s, $CH_3$a), 1.35 (3H, s, $CH_3$b); $\delta_C$ (125 MHz, DMSO-d6): 156.8 (C2Ar), 156.0 (C4Ar), 152.6 (C2'Ar), 149.3 (C8Ar), 144.2 (C5'Ar), 140.2 (C6Ar), 177.9 (C9Ar), 113.0 (Acetyl C), 112.0 (C3'Ar), 111.4 (C4'Ar), 89.0 (C2Furyl), 86.8 (C5Furyl), 83.3 (C3Furyl), 81.5 (C4Furyl), 61.6 ($CH_2$), 27.9 ($CH_3$a), 25.2 ($CH_3$b); $\nu_{max}$/ $cm^{-1}$ (solid): 3419, 3311, 2987, 2938, 1639, 1547, 1369; HPLC: $T_r$= 2.20 (57% rel. area); *m/z* (ES): (Found: $[M+H]^+$, 374.1462. $C_{17}H_{19}N_5O_5$ requires *[M+H]*, 374.1459. $[\alpha]_D$ = -44.8° (c 0.1, MeOH).

**Preperation of [(3aR, 4R, 6R, 6aR)-6-(6-amino-2-(furan-2-yl)-9H-purin-9-yl)-2,2-dimethyl-tetrahydro-2H-furo[3,4-d][1,3]dioxol-4-yl]methyl sulfamate**

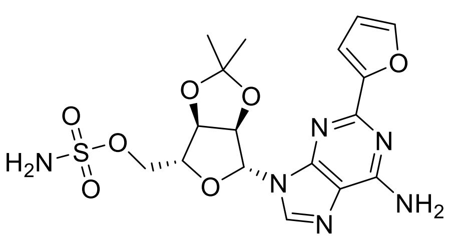


Preparation was *via* general method B using 2'3'-O-Isopropylidene-2-(furan-2-yl)adenosine (0.25g, 0.67 mmol) to afford the desired product as a colourless glassy solid (0.20 g, 0.45 mmol, 67%) m.p.: 83.3-85.2 °C; *R*f: 0.46 (9:1 Chloroform-MeOH); $\delta_H$ (500 MHz, DMSO-d6): 8.31 (1H, s, 6-HAr), 7.83 (1H, s, 3'-HAr), 7.56 (2H, brs, $SNH_2$), 7.47 (2H, brs, $NH_2$), 7.12 (1H, d, J 3.2, 5'-HAr), 6.65 (1H, dd, J 3.2 and 1.8, 4'-HAr), 6.30 (1H, d, J 1.9, 2-HFuryl), 5.44 (1H, d, J 6.2 3-HFuryl), 5.23 (1H, dd, J 6.2 and 3.5, 4-HFuryl), 4.43 (1H, app s, 5-HFuryl), 4.35-4.15 (2H, m, $CH_2$), 1.58 (3H, s, $CH_3$a), 1.36 (3H, s, $CH_3$b); $\delta_C$ (125 MHz, DMSO-d6): 159.5 (C2Ar), 155.8 (C4Ar), 152.3 (C1'Ar), 150.6 (C8Ar), 145.0 (C3'Ar), 140.0 (C6Ar), 119.4 (C9Ar), 114.4 (Acetyl C), 113.3 (C5'Ar), 112.1 (C4'Ar), 90.7 (C2Furyl), 84.4 (C5Furyl), 83.3 (C3Furyl), 81.1 (C4Furyl), 70.5 ($CH_2$), 26.7 ($CH_3$a), 24.5 ($CH_3$b); $\nu_{max}$/ $cm^{-1}$ (solid): 3319, 3161, 2988, 1628, 1577, 1361; HPLC: $T_r$= 2.27 (51% rel. area); *m/z* (ES): (Found: $[M+H]^+$, 453.1189. $C_{17}H_{20}N_6O_7S$ requires *[M+H]*, 453.1187. $[\alpha]_D$ = 1.4° (c 0.1, MeOH).

**Preparation of tert-butyl N-[(2S)-1-[((([(3aR, 4R, 6R, 6aR)-6-(6-amino-2-(furan-2-yl)-9H-purin-9-yl)-2,2-dimethyl-tetrahydro-2H-furo[3,4-d][1,3]dioxol-4-yl]methoxy)sulfonyl)amino]-3-(benzyloxy)-1-oxopropan-2-yl]carbamate**

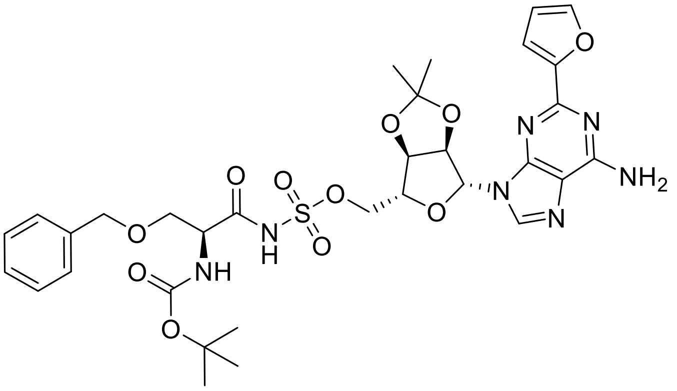


Preparation was *via* general method C using [(3aR, 4R, 6R, 6aR)-6-(6-amino-2-(furan-2-yl)-9H-purin-9-yl)-2,2-dimethyl-tetrahydro-2H-furo[3,4-d][1,3]dioxol-4-yl]methyl sulfamate (0.20 g, 0.44 mmol) to afford the desired product as a colourless glassy solid (0.10 g, 0.14 mmol, 32%) m.p.:157.7-159.4 °C; *R*f: 0.50 (9:1 DCM-

MeOH); $\delta_H$ (500 MHz, DMSO-d6): 8.40 (1H, s, 6-HAr), 7.81 (1h, s, 3'-HAr), 7.42 (2H, brs, NH2), 7.32-7.28 (5H, m, Bzl), 7.24 (1H, d, J 4.1, 5'-HAr), 6.62 (1H, dd, J 4.1 and 1.87, 4'-HAr), 6.21 (1H, d, J 2.8, 2-HFuryl), 6.06 (1H, app s, NH), 5.36 (1H, app s, 3-HFuryl), 5.08 (1H, app s, 4-HFuryl), 4.48-4.32 (3H, m, $CH_2$ Bzl and 5-HFuryl), 4.15-3.90 (3H, m, $CH_2O$ and Chiral H), 3.70-3.50 (2H, m, $CH_2$Chiral), 1.57 (3H, s, $CH_3$a), 1.38 (9H, s, $CH_3 \times 3$), 1.33 (3H, s, $CH_3$b); $\delta_C$ (125 MHz, DMSO-d6): 170.0 (C=ONH), 158.3 (C2Ar), 156.4 (C4Ar), 155.3 (C=OBOC), 152.5 (C1Furan), 150.6 (C8Ar), 144.7 (C3Furan), 140.2 (C6Ar), 137.7 (C1Bzl), 128.8-127.7 (Aromatics), 119.3 (C9Ar), 114.2 (Acetyl C), 112.4 (C5Furan), 111.9 (C4Furan), 89.2 (C2Furyl), 84.4 (C5Furyl), 84.1 (C3Furyl), 82.1 (C4Furyl), 79.6 (BOC C), 72.2 (CH2 Bzl), 71.0 (CH2O), 67.7 (CH2 Chiral), 56.9 (Chiral C), 28.7 (CH3 × 3), 27.6 (CH3a), 25.7 (CH3b); $\nu_{max}$/ $cm^{-1}$ (solid): 3342, 1674, 1572, 1364; HPLC: $T_r$= 2.51 (73% rel. area); *m/z* (ES): (Found: $[M+H]^+$, 730.2513. $C_{32}H_{39}N_7O_{11}S$ requires *[M+H]*, 730.2501. $[\alpha]_D$ = -24.8° (c 0.1, MeOH).

**Preparation of (2S)-2-amino-1-[([[(2R, 3S, 4R, 5R)-5-(6-amino-2-(furan-2-yl)-9H-purin-9-yl)-3,4-dihydroxyoxolan-2-yl]methoxy)sulfonyl)amino]-3-hydroxypropan-1-one (6)**

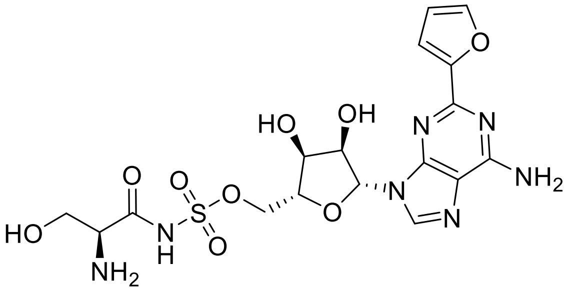


Preparation was via general method D using tert-butyl N-[(2S)-1-[(([(3aR, 4R, 6R, 6aR)-6-(6-amino-2-(furan-2-yl)-9H-purin-9-yl)-2,2-dimethyl-tetrahydro-2H-furo[3,4-d][1,3]dioxol-4-yl]methoxy)sulfonyl)amino]-3-(benzyloxy)-1-oxopropan-2-yl]carbamate (100 mg, 0.14 mmol) to afford the desired product as a colourless powder (14.6 mg, 0.03 mmol, 21%) m.p.: 117.2 (Decomp); $R_f$: Baseline (9:1 DCM-MeOH); $\delta_H$ (500 MHz, DMSO-d6): 8.54 (1H, brs, NH), 8.34 (2H, brs, $NH_2$), 8.14 (1H, s, 6-HAr), 7.96 (1H, app s, 3'-HAr), 7.46 (1H, app t, J 10.1, 5'-HAr), 6.76 (1H, dd, J 10.1 and 8.5, 4'-HAr), 6.02 (1H, d, J 5.2, 2-HFuryl), 4.65 (1H, d, J 5.2, 3-HFuryl), 4.52-4.45 (2H, m, $CH_2O$), 4.29 (1H, app s, 4-HFuryl), 4.20 (1H, app s, 5-HFuryl), 3.89-3.75 (3H, m, $CH_2$ Chiral and Chiral H); $\delta_C$ (125 MHz, DMSO-d6): 172.0 (C=O), 158.1 (C2Ar), 156.1 (C4Ar), 152.5 (C1'Ar), 150.6 (C8Ar), 145.3 (C3'Ar), 141.2 (C6Ar), 117.9 (C9Ar), 113.9 (C5'Ar), 113.2 (C4'Ar), 88.2 (C2Furyl), 84.6 (C5furyl), 73.4 (C3Furyl), 71.8 (C4Furyl), 60.6 ($CH_2O$), 54.9 (Chiral C), 45.2 ($CH_2$ Chiral); $\nu_{max}$/ $cm^{-1}$ (solid): 3089, 1689, 1595, 1477, 1383; *m/z* (ES): (Found: $[M+H]^+$,. $C_{17}H_{21}N_7O_9S$ requires *[M+H]*, 500.1194.) $[\alpha]_D$ = 22.9° (c 0.1, MeOH).

**Preparation of 2'3'-O-Isopropylidene-2-(thiophen-3-yl)adenosine**

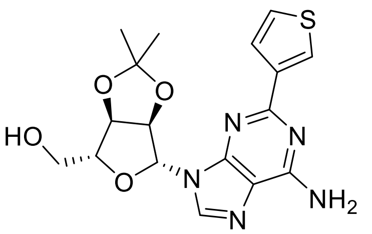

Preparation was via general method A using 2'3'-O-Isopropylidene-2-chloroadenosine (0.50 g, 1.46 mmol) and 3-thienyl boronic acid (0.75 g, 5.85 mmol) to afford the desired product as a colourless powder (0.46 g, 1.17 mmol, 80%); m.p.: 100.1-102.0 °C; *R*f: 0.62 (9:1 Chloroform-MeOH); $\delta_H$ (500 MHz, DMSO-d6): 8.34 (1H, s, 6-HAr), 8.18 (1H, dd, J 3.1 and 1.2, 2'-HAr), 7.79 (1H, dd, J 5.0 and 1.2, 4'-HAr), 7.42 (1H, dd, J 5.0 and 3.1, 5'-H), 7.36 (2H, brs, $NH_2$), 6.23 (1H, d, J 6.2, 2-HFuryl), 5.49 (1H, dd, J 6.2 and 2.7, 3-HFuryl), 5.11 (1H, dd, J 6.2 and 2.7, 4-HFuryl), 4.22 (1H, t, J 5.5 and 2.7, 5-HFuryl), 3.59 (2H, dd, J 11.5 and 5.5, $CH_2$), 1.57 (3H, s, $CH_3$a), 1.36 (3H, s, $CH_3$b); $\delta_C$ (125 MHz, DMSO-d6): 173.2 (C2Ar), 155.8 (C4Ar), 140.3 (C6Ar), 127.4 (C4'Ar), 125.1 (C5'Ar), 120.0 (C2'Ar), 119.2 (C9Ar), 116.7 (C1'Ar), 114.2 (AcetylC), 90.6 (C2Furyl), 87.3 (C5Furyl), 83.6 (C3Furyl), 81.2 (C4Furyl), 61.8 ($CH_2$), 26.7 ($CH_3$a), 25.0 ($CH_3$b); $\nu_{max}$/ $cm^{-1}$ (solid): 3253, 2939, 1633, 1586, 1344; HPLC: $T_r$= 2.19 (79% rel. area); *m/z* (ES): (Found: $[M+H]^+$, 365.1058. $C_{17}H_{19}N_5O_4S$ requires *[M+H]*, 365.1051. $[\alpha]_D$ = -6.6° (c 0.1, MeOH).

**Preperation of [(3aR, 4R, 6R, 6aR)-6-(6-amino-2-(thiophen-3-yl)-9H-purin-9-yl)-2,2-dimethyl-tetrahydro-2H-furo[3,4-d][1,3]dioxol-4-yl]methyl sulfamate**

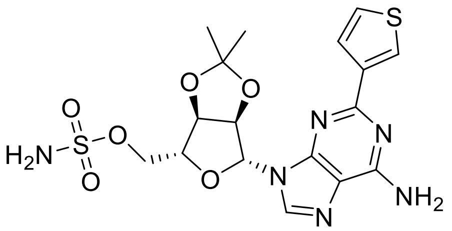

Preparation was *via* general method B using 2'3'-O-Isopropylidene-2-(thiophen-3-yl)adenosine (0.45 g, 1.16 mmol) to afford the desired product as a pale brown oil (0.51 g, 1.09 mmol, 94%) *R*f: 0.48 (9:1 Chloroform-MeOH); $\delta_H$ (500 MHz, DMSO-d6): 8.31 91H, s, 6-HAr), 8.18 (1H, d, J 2.0, 2'-HAr), 7.77 (1H, dd, J 6.1 and 2.0, 5'-HAr), 7.58 (2H, brs, $SNH_2$), 7.42 (1H, d, J 6.1, 4'-H), 7.38 (2H, brs, $NH_2$), 6.31 (1H, d, J 2.2, 2-HFuryl), 5.54 (1H, d, J 4.0, 3-HFuryl), 5.20 (1H, d, J 4.0, 4-H Furyl), 4.44 (1H, app s, 5-HFuryl), 4.35-4.15 (2H, m, $CH_2$), 1.60 (3H, s, $CH_3$a), 1.37 (3H, s, $CH_3$b); $\delta_C$ (125 MHz, DMSO-d6): 169.5 C2Ar), 155.9 (C4Ar), 149.4 (C8Ar), 142.2 (C6Ar), 134.8 (C4'Ar), 132.4 (C5'Ar), 126.4 (C9Ar), 125.0 (C2'Ar), 117.9 (C1'Ar), 113.5 (Acetyl C), 88.9 (C2Furyl), 83.7 (C5Furyl), 83.2 (C3Furyl), 81.2 (C4Furyl), 68.1 ($CH_2$), 26.7 ($CH_3$a), 25.1 ($CH_3$b); $\nu_{max}$/ $cm^{-1}$ (solid): 3324, 3200, 2988, 1619, 1398; HPLC: $T_r$= 2.23 (82% rel. area); *m/z* (ES): (Found: $[M+H]^+$, 469.0958. $C_{17}H_{20}N_6O_6S_2$ requires *[M+H]*, 469.0959. $[\alpha]_D$ = 6.0° (c 0.1, MeOH).

**Preparation of tert-butyl N-[(2S)-1-[(([(3aR, 4R, 6R, 6aR)-6-(6-amino-2-(thiophen-3-yl)-9H-purin-9-yl)-2,2-dimethyl-tetrahydro-2H-furo[3,4-d][1,3]dioxol-4-yl]methoxy)sulfonyl)amino]-3-(benzyloxy)-1-oxopropan-2-yl]carbamate**

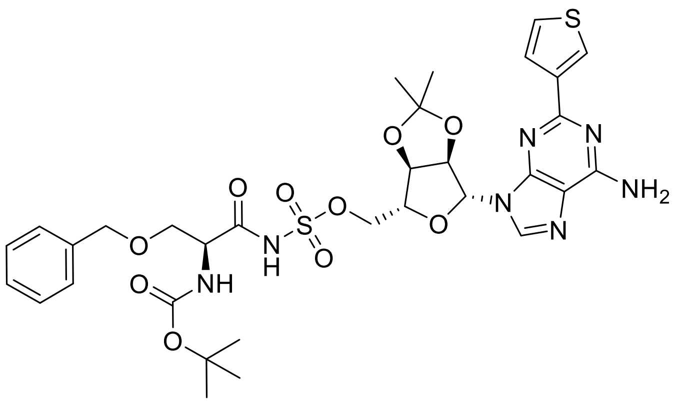

Preparation was *via* general method C using [(3aR, 4R, 6R, 6aR)-6-(6-amino-2-(thiophen-3-yl)-9H-purin-9-yl)-2,2-dimethyl-tetrahydro-2H-furo[3,4-d][1,3]dioxol-4-yl]methyl sulfamate (0.50 g, 1.06 mmol) to afford the desired product as a colourless glassy solid (0.33 g, 0.45 mmol, 42%) m.p.: 156.2-158.0 °C; $R_f$: 0.58 (9:1 DCM-MeOH); $\delta_H$ (500 MHz, DMSO-d6): 8.38 (1H, s, 6-HAr), 8.19 (1H, dd, J 3.1 and 1.1, 2'-HAr), 7.78 (1h, dd, J 5.1 and 1.1 4'-HAr), 7.57 (1H, dd, J 5.0 and 3.1, 5'-HAr), 7.30 (2H, brs, $NH_2$), 7.29-7.25 (5H, m, bzl), 6.23 (1H, d, J 3.0, 2-HFuryl), 6.07 (1H, app s, NH), 5.41 (1H, d, J 2.8, 3-HFuryl), 5.07 (1H, d, J 2.8, 4-HFuryl), 4.41 (3H, app d, J 9.9, $CH_2$Bzl) and 5-HFuryl, 4.10-4.00 (2H, m, $CH_2O$), 3.98-3.94 (1H, m, Chiral H), 3.70-3.50 (2H, m, $CH_2$Chiral), 1.58 (3H, s, $CH_3$a), 1.39 (9H, s, $CH_3$ × 3), 1.35 (3H, s, $CH_3$b); $\delta_C$ (125 MHz, DMSO-d6): 173.2 (C2Ar), 170.1 (C=ONH), 156.1 (C4Ar), 155.9 (C=OBOC), 150.8 (C8Ar), 142.8 (C6Ar), 138.5 (C1Bzl), 128.0 (C5Bzl and C3Bzl), 127.3 (C6Bzl and C2Bzl), 127.1 (C4Bzl), 126.1 (C4Thio), 125.1 (C5Thio), 120.0 (C2Thio), 119.4 (C9Ar), 117.0 (C1Thio), 113.1 (Acetyl C), 88.7 (C2Furyl), 83.6 (C5Furyl), 83.3 (C3Furyl), 81.6 (C4Furyl), 79.7 (BOC C), 73.4 ($CH_2$ Bzl), 71.7 ($CH_2O$), 68.9 ($CH_2$ Chiral), 56.2 (Chiral C), 28.2 ($CH_3$ × 3), 27.1 ($CH_3$a), 25.2 ($CH_3$b); $\nu_{max}$/ $cm^{-1}$ (solid): 3250, 3052, 1675, 1526, 1374; HPLC: $T_r$= 2.53 (100% rel. area); *m/z* (ES): (Found: $[M+H]^+$, 746.2286. $C_{32}H_{39}N_7O_{10}S_2$ requires *[M+H]*, 746.2273. $[\alpha]_D$ = 28.9° (c 0.1, MeOH).

**Preparation of (2S)-2-amino-1-[([[(2R, 3S, 4R, 5R)-5-(6-amino-2-(thiophen-3-yl)-9H-purin-9-yl)-3,4-dihydroxyoxolan-2-yl]methoxy)sulfonyl)amino]-3-hydroxypropan-1-one (7)**

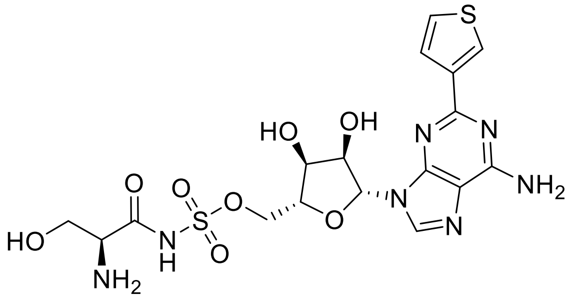

Preparation was via general method D using tert-butyl N-[(2S)-1-[(([(3aR, 4R, 6R, 6aR)-6-(6-amino-2-(thiophen-3-yl)-9H-purin-9-yl)-2,2-dimethyl-tetrahydro-2H-furo[3,4-d][1,3]dioxol-4-yl]methoxy)sulfonyl)amino]-3-(benzyloxy)-1-oxopropan-2-yl]carbamate (0.30 g, 0.40 mmol) to afford the desired product as a colourless powder (27.5 mg, 0.05 mmol, 13%) m.p.: 108.3 °C (Decomp); $R_f$: Baseline (9:1 DCM-MeOH); $\delta_H$ (500 MHz,

DMSO-d6): 8.46 (1H, brs, NH), 8.34 (1H, s, 2'-HAr), 8.14 (1H, s, 6-HAr), 8.00 (2H, brs, NH2), 7.87-7.78 (1H, m, 4'-HAr), 7.68-7.59 (1H, m, 5'-HAr), 6.00 (1H, d, J 5.5, 2-HFuryl), 4.70 (1H, t, J 5.5, 3-HFuryl), 4.37-4.22 (4H, m, $CH_2O$, 4-HFuryl and 5-HFuryl), 3.83-3.65 (3H, m, $CH_2$ chiral and chiral H); $\delta_C$ (125 MHz, DMSO-d6): 172.0 (C=O), 163.5 (C2Ar), 156.0 C4Ar), 150.7 C8Ar), 140.1 (C6Ar), 127.8 (C4'Ar), 127.1 (C5'Ar), 120.0 (C2'Ar), 119.2 (C9Ar), 116.8 (C1'Ar), 88.2 (C2Furyl), 84.2 (C5furyl), 73.7 (C3Furyl), 71.8 (C4Furyl), 60.8 ($CH_2O$), 57.2 (Chiral C), 45.4 ($CH_2$ chiral); $\nu_{max}$/ $cm^{-1}$ (solid): 3096, 1685, 1588, 1402, 1275; *m/z* (ES): (Found: $[M+H]^+$, 516.0972. $C_{17}H_{21}N_7O_8S_2$ requires *[M+H]*, 516.0966.) $[\alpha]_D$ = 20.7° (c 0.1, MeOH).

**Preparation of 2'3'-O-Isopropylidene-2-(pyridine-3-yl)adenosine**

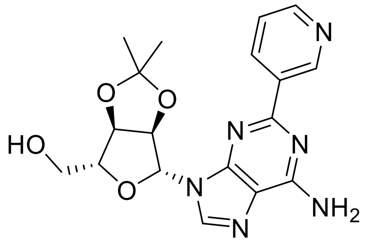


Preparation was via general method A using 2'3'-O-Isopropylidene-2-chloroadenosine (0.50 g, 1.46 mmol) and 3-pyridine boronic acid (0.71 g, 5.85 mmol) to afford the desired product as a colourless powder (0.29 g, 0.75 mmol, 51%); m.p.: 122.4-124.2 °C; $R_f$: 0.53 (9:1 Chloroform-MeOH); $\delta_H$ (500 MHz, DMSO-d6): 9.49 (1h, s, 2'-HAr), 8.67-8.61 (2H, m, 6'-HAr and 4'-HAr), 8.40 (1H, s, 6-HAr), 7.52 (3H, brs, $NH_2$ and 5'-HAr), 6.26 (1H, d, J 2.7, 2-HFuryl), 5.52 (1H, dd, J 6.1 and 2.7, 3-HFuryl), 5.10 (1H, dd, J 6.1 and 2.9, 4-HFuryl), 5.05 (1H, t, J 5.5, OH), 4.23 (1H, dd, J 8.2 and 5.3, 5-HFuryl), 3.62-3.51 2H, m, $CH_2$), 1.57 (3H, s, $CH_3$a), 1.36 (3H, s, $CH_3$b); $\delta_C$ (125 MHz, DMSO-d6): 160.3 (C2Ar), 156.5 (C4Ar), 150.8 (C4Pip), 150.2 (C8Ar), 149.5 (C2Pip), 141.1 (C6Ar), 134.5 (C6Pip), 134.1 (C1Pip), 124.0 (C5Pip), 119.0 (C9Ar), 113.5 (Acetyl C), 89.7 (C2Furyl), 87.2 (C5Furyl), 83.8 (C3furyl), 81.9 (C4Furyl), 62.0 ($CH_2$), 27.6 ($CH_3$a), 25.7 ($CH_3$b); $\nu_{max}$/ $cm^{-1}$ (solid): 3322, 1632, 1572, 1375; HPLC: $T_r$= 1.99 (100% rel. area); *m/z* (ES): (Found: $[M+H]^+$, 385.1626. $C_{18}H_{20}N_6O_4$ requires *[M+H]*, 385.1619. $[\alpha]_D$ = -1.4° (c 0.1, MeOH).

**Preperation of [(3aR, 4R, 6R, 6aR)-6-(6-amino-2-(pyridine-3-yl)-9H-purin-9-yl)-2,2-dimethyl-tetrahydro-2H-furo[3,4-d][1,3]dioxol-4-yl]methyl sulfamate**

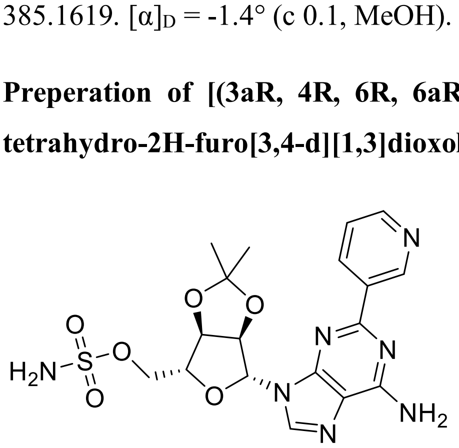


Preparation was *via* general method B using 2'3'-O-Isopropylidene-2-(pyridine-3-yl)adenosine (0.29 g, 0.75 mmol) to afford the desired product as a colourless glassy solid ( 0.25 g, 0.54 mmol, 72%) m.p.: 93.1-94.7 °C; $R_f$: 0.39 (9:1 Chloroform-MeOH); $\delta_H$ (500 MHz, DMSO-d6):9.48 (1H, s, 2'-HAr), 8.67-8.59 (2H, m, 6'-H and 4'-H), 8.37 (1h, s, 6-HAr), 7.65-7.50 (5H, m, $SNH_2$, $NH_2$ and 3'-HAr), 6.34 (1H, d, J 2.2, 2-HFuryl), 5.58 (1H, d, J 3.7 3-HFuryl), 5.19 (1H, dd, J 6.0 and 3.7, 4-HFuryl), 4.45 (1H, app s, 5-HFuryl), 4.30-4.20 (2H, m, $CH_2$), 1.60 (3H, s, $CH_3$a), 1.39 (3H, s, $CH_3$b) ; $\delta_C$ (125 MHz, DMSO-d6): 161.7 (C2Ar), 155.8 C4Ar), 150.8 (C4'Ar), 150.6 (C8Ar), 149.5 (C2'Ar), 141.1

(C6Ar), 135.4 (C6'Ar), 133.2 (C1'Ar), 124.0 (C5'Ar), 119.1 (C9Ar), 114.2 (Acetyl C), 89.4 (C2Furyl), 84.0 (C5Furyl), 83.7 (C3Furyl), 81.6 (C4Furyl), 68.0 ($CH_2$), 27.5 ($CH_3$a), 25.7 ($CH_3$b); $\nu_{max}$/ $cm^{-1}$ (solid): 3345, 3180, 2985, 1632, 1580, 1374; HPLC: $T_r$= 2.04 (100% rel. area); *m/z* (ES): (Found: $[M+H]^+$, 464.1357. $C_{18}H_{21}N_7O_6S$ requires *[M+H]*, 464.1347. $[\alpha]_D$ = 53.5° (c 0.1, MeOH).

**Preparation of tert-butyl N-[(2S)-1-[(([(3aR, 4R, 6R, 6aR)-6-(6-amino-2-(pyridine-3-yl)-9H-purin-9-yl)-2,2-dimethyl-tetrahydro-2H-furo[3,4-d][1,3]dioxol-4-yl]methoxy)[6]sulfonyl)amino]-3-(benzyloxy)-1-oxopropan-2-yl]carbamate**

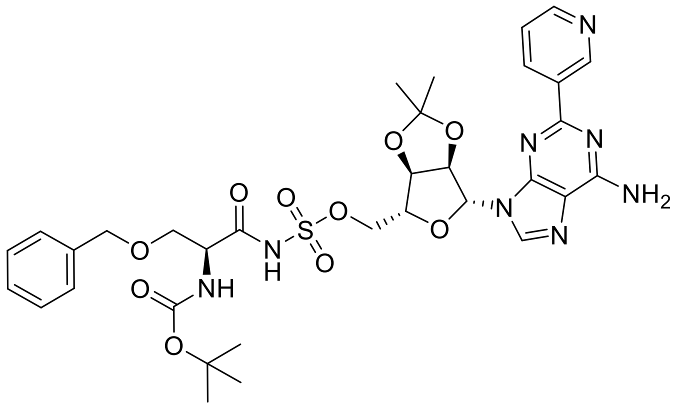


Preparation was *via* general method C using [(3aR, 4R, 6R, 6aR)-6-(6-amino-2-(pyridine-3-yl)-9H-purin-9-yl)-2,2-dimethyl-tetrahydro-2H-furo[3,4-d][1,3]dioxol-4-yl]methyl sulfamate (0.25 g, 0.54 mmol) to afford the desired product as a colourless glassy solid (0.12 g, 0.16 mmol, 29%) m.p.: 167.2-164.5 °C; $R_f$: 0.39 (9:1 DCM-MeOH); $\delta_H$ (500 MHz, DMSO-d6): 9.49 (1H, s, 2'-HAr), 8.64 (2H, d, J 5.7, 6'-HAr and 4'-HAr), 8.46 (1H, s, 6-HAr), 7.52 (3H, brs, $NH_2$ and 5'-HAr), 7.27-7.24 (5H, m, Bzl), 6.27 (1H, d, J 3.0, 2-HFuryl), 6.08 (1H, d, J 7.8, NH), 5.43 (1H, d, J 2.7, 3-HFuryl), 5.07 (1H, app s, 4-Hfuryl), 4.41 (3H, app, s, $CH_2$ Bzl and 5-HFuryl), 4.05 (2H, d, J 4.6, $CH_2O$), 3.94 (1H, s, Chiral H), 3.62 (2H, d J 6.6, $CH_2$ Chiral), 1.59 (3H, s, $CH_3$a), 1.36 (9H, s, $CH_3$ × 3), 1.35 (3H, s, $CH_3$b); $\delta_C$ (125 MHz, DMSO-d6): 173.7 (C=ONH), 160.5 (C2Ar), 156.1 (C4Ar), 155.4 (C=OBOC), 150.2 (C4Pip), 150.1 (C8Ar), 148.9 (C2Pip), 140.1 (C6Ar), 137.7 (C1Bzl), 134.9 (C6Pip), 133.2 (C1Pip), 128.0 (C5Bzl and C3Bzl), 127.3 (C6Bzl and C2Bzl), 127.1 (C4Bzl), 123.5 (C5Pip), 119.4 (C9Ar), 114.2 (Acetyl C), 89.0 (C2Furyl), 83.5 (C5Furyl), 83.4 (C3Furyl), 81.5 (C4Furyl), 79.6 (BOC C), 73.7 ($CH_2$ Bzl), 71.7 ($CH_2O$), 68.2 ($CH_2$ Chiral), 54.7 (Chiral C), 28.2 ($CH_3$ × 3), 27.1 ($CH_3$a), 25.2 ($CH_3$b); $\nu_{max}$/ $cm^{-1}$ (solid): 3377, 2976, 1662, 1573, 1369; HPLC: $T_r$= 2.37 (100% rel. area); *m/z* (ES): (Found: $[M+H]^+$, 741.2674. $C_{33}H_{40}N_8O_{10}S$ requires *[M+H]*, 741.2661. $[\alpha]_D$ = -13.6° (c 0.1, MeOH).

**Preparation of (2S)-2-amino-1-[([[(2R, 3S, 4R, 5R)-5-(6-amino-2-(pyridine-3-yl)-9H-purin-9-yl)-3,4-dihydroxyoxolan-2-yl]methoxy)sulfonyl)amino]-3-hydroxypropan-1-one (8)**

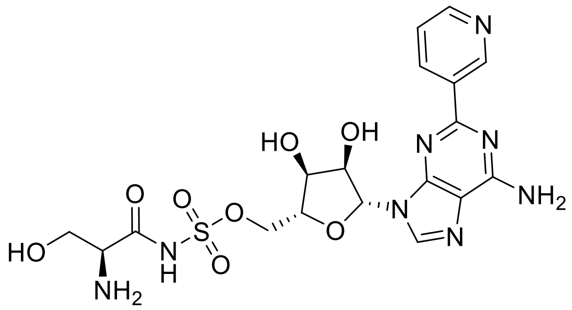

Preparation was via general method D using tert-butyl N-[(2S)-1-[(([(3aR, 4R, 6R, 6aR)-6-(6-amino-2-(pyridine-3-yl)-9H-purin-9-yl)-2,2-dimethyl-tetrahydro-2H-furo[3,4-d][1,3]dioxol-4-yl]methoxy)sulfonyl)amino]-3-(benzyloxy)-1-oxopropan-2-yl]carbamate (100 mg, 0.13 mmol) to afford the desired product as a colourless powder (23.8 mg, 0.05 mmol, 36%) m.p.: 120.2 °C (Decomp); $R_f$: Baseline (9:1 DCM-MeOH); $\delta_H$ (500 MHz, DMSO-d6): 9.50 (1H, s, 2'-HAr), 8.95-8.75 (2H, m, 6'-HAr and 4'-HAr), 8.47 (1H, s, 6-HAr), 7.93 (2H, brs, $NH_2$), 7.54-7.50 (1H, m, 5'-HAr), 6.04 (1H, d, J 6.0 2-HFuryl), 4.82-4.72 (1H, m, 3-HFuryl), 4.37-4.23 2H, m, $CH_2$*O and 4-HFuryl), 4.18 (2H, app d, J 8.9, 5-HFuryl and CH2*O), 3.82 (1H, d, J 8.0 $CH_2$ chiral), 3.73-3.60 (1H, m, $CH_2$ chiral), 3.58-3.54 (1H, m, Chiral H); $\delta_C$ (125 MHz, DMSO-d6): 171.9 (C=O), 160.3 (C2Ar), 156.1 (C4Ar), 151.0 (C4'Ar), 150.8 (C8Ar), 150.7 (C2'Ar), 141.2 (C6Ar), 138.6 (C2'Ar), 135.6 (C6'Ar), 133.4 (C1'Ar), 125.2 (C5'Ar), 119.4 (C9Ar), 87.9 (C2Furyl), 82.9 (C5Furyl), 73.7 (C3Furyl), 71.2 (C4Furyl), 68.7 ($CH_2O$), 60.8 ($CH_2$ chiral), 57.5 (Chiral C); $\nu_{max}$/ $cm^{-1}$ (solid): 3317, 3118, 1633, 1587, 1587, 1382; *m/z* (ES): (Found: $[M+H]^+$, 511.1356. $C_{18}H_{22}N_8O_8S$ requires *[M+H]*, 511.1356. $[\alpha]_D$ = 19.3° (c 0.1, MeOH).
